# Edge-based growth control in *Arabidopsis* involves two cell wall-associated Receptor-Like Proteins

**DOI:** 10.1101/2022.06.10.495700

**Authors:** Liam Elliott, Monika Kalde, Ann-Kathrin Schuerholz, Sebastian Wolf, Ian Moore, Charlotte Kirchhelle

## Abstract

Morphogenesis of multicellular organs requires coordination of cellular growth. In plants, 3D growth is driven by undirected turgor pressure, whereas growth directionality is controlled by cell wall mechanical properties at 2D cell faces. Their shared cell wall also fixes cells in their position, and plants thus have to integrate tissue-scale mechanical stresses arising due to growth in a fixed tissue topology. This implies a need to monitor cell wall mechanical and biochemical status and to adapt growth accordingly. Here, we propose that plant cells use their 1D cell edges to monitor cell wall status. We describe two Receptor-Like Proteins, RLP4 and RLP4-L1, which occupy a unique polarity domain at cell edges established through a targeted secretory transport pathway. We show that at cell edges, RLP4s associate with the cell wall via their extracellular domain, and contribute to directional growth control in *Arabidopsis*.

## Introduction

Organs of multicellular organisms arise through cellular growth: the irreversible increase in volume of each cell within the organ. To develop a defined organ shape, neighbouring cells need to coordinate their growth. This can occur through tissue-scale organising cues (morphogen gradients or stress fields), but at the local scale, heterogeneities in cellular growth can still cause mechanical conflict. In animal systems, such local conflicts can be relaxed through changes in tissue topology. In plants however, neighbouring cells are surrounded by a shared cell wall and cannot move relative to each other. Within the confines of this fixed tissue topology, mechanical conflicts have to be otherwise accommodated.

Plant cell growth is driven by non-directional turgor pressure, which is translated into directional growth through construction and modification of a cell wall with heterogeneous biochemical and mechanical properties (Cosgrove, 2018). Cell walls consist of water, carbohydrates (cellulose, hemicelluloses, and pectins), and a small fraction of proteins (Carpita and Gibeaut, 1993). Plants primarily control growth direction through oriented deposition of cellulose fibres of high tensile strength, which constrain growth parallel to their net orientation (Green, 1964) and are locally reinforced through interactions with hemicelluloses (Cosgrove, 2014). Pectins influence cell wall porosity, but can also contribute to differential extensibility of the cell wall (Haas et al., 2020; Lin et al., 2022; Peaucelle et al., 2015). Despite their distinct structures and mechanical properties, loss of specific cell wall components can be compensated by others: for example, pectins assume a more prominent role as a load-bearing component in plant cell walls lacking the hemicellulose xyloglucan (Park and Cosgrove, 2012). This implies that plant cells can perceive changes in their cell wall status and adapt their cell wall biogenesis accordingly.

Several cell surface receptors families, including Wall-Associated Kinases (WAKs), *Catharanthus roseus* Receptor-Like Kinase 1-Likes (CrRLK1Ls) and Receptor-Like Proteins (RLPs), have been linked to perception of cell wall integrity or responses to depletion of cell wall components (Boisson-Dernier et al., 2009; Decreux and Messiaen, 2005; Deslauriers and Larsen, 2010; Feng et al., 2018; Guo et al., 2009; Hematy et al., 2007; Kohorn et al., 2009; Lally et al., 2001; Wolf et al., 2014). Some of these cell surface receptors can directly interact with cell wall carbohydrates – for example, WAKs can bind to pectins *in vivo* and *in vitro* (Kohorn, 2016). CrRLK1L family members including the intensively-studied FERONIA (FER) carry a Malectin-like Domain (MLD) in their extracellular region, which can bind to pectin *in vitro* and *in vivo* (Feng et al., 2018; Lin et al., 2022; Tang et al., 2022). FER can also indirectly interact with the cell wall through interaction with cell-wall-bound Leucine-Rich Repeat Extensins (LRXs), an interaction that has been functionally linked to cell expansion (Dünser et al., 2019). Signalling involving various CrRLK1Ls including FER also depends on binding of RAPID ALKALINAZATION FACTOR (RALF) peptides (Ge et al., 2017; Gonneau et al., 2018; Haruta et al., 2014). In other cases, receptors have been functionally linked to cell wall sensing even though their ligands have not been identified. For example, the Receptor-Like Protein RLP44 is involved in mediating cellular responses to changes in the pectin methyl-esterification status through co-opting the Brassinosteroid (BR) signalling pathway via direct interaction with two RLKs involved in BR signalling, BRASSINOSTEROID INSENSITIVE 1 (BRI1) and BRI1-ASSOCIATED KINASE 1 ((BAK1; (Holzwart et al., 2018, 2020; Wolf et al., 2014)). Despite the identification of such ligands and interacting partners for several cell surface receptors, the role of these cell wall sensing systems in the assembly and continuous modification of the cell wall required during normal growth is not well understood. One reason for this may be a lack of appreciation of the spatial context (i.e., the 3D geometry of the cell) in which such signals are perceived and translated into cell wall assembly and growth.

Here, we describe two Receptor-Like Proteins (RLP4s), that occupy a unique subcellular domain in the plasma membrane (PM) of growing cells: the geometric edges (where two faces of a polyhedral cell meet). We show that RLP4s localisation at PM edges is established through a secretory pathway specifically targeted to cell edges mediated by the small GTPase RAB-A5c. We also show that at the cell surface, RLP4s associate with the cell wall, and act as core components of an edge-based growth control system in *Arabidopsis* lateral roots.

## Results

### Two Receptor-Like Proteins are interactors of the edge-localised small GTPase RAB-A5c

We have previously shown that the plant-specific GTPase RAB-A5c mediates a secretory pathway from the trans-Golgi network/early endosome (TGN/EE) to the edges of plant cells, where two cell faces meet (Kirchhelle et al., 2016). To identify interactors of RAB-A5c, we used a comparative proteomics approach: we performed co-immunoprecipitation coupled with label-free semi-quantitative mass spectrometry (co-IP-MS) against YFP:RAB-A5c (Kirchhelle et al., 2016) and two related Rab GTPases: the late endosome/tonoplast-localised YFP:RAB-G3f (Geldner et al., 2009) and the TGN/EE-localised YFP:RAB-A2a (Chow et al., 2008). We then identified proteins significantly enriched in the YFP:RAB-A5c interactome compared to the interactomes of YFP:RAB-G3f and YFP:RAB-A2a (Figure 1A-C) to distinguish generic interactors of Rab GTPases or of the Rab-A subfamily from those specific to RAB-A5c. Our pipeline identified 120 proteins significantly enriched in the YFP:RAB-A5c interactome compared to both YFP:RAB-G3f and YFP:RAB-A2a, which we subsequently ranked based on their relative enrichment as well as overall abundance in the YFP:RAB-A5c interactome while penalising high abundance in a Col-0 negative control (supplementary table 1). Within the top twenty candidates identified in this approach, we found two related proteins: RECEPTOR-LIKE PROTEIN 4 (RLP4; (Wang et al., 2008)) and its closest relative in *Arabidopsis*, At1g25570, which we will refer to as RECEPTOR-LIKE PROTEIN 4-LIKE1 (RLP4-L1; Figure 1D).

**Figure 1:**
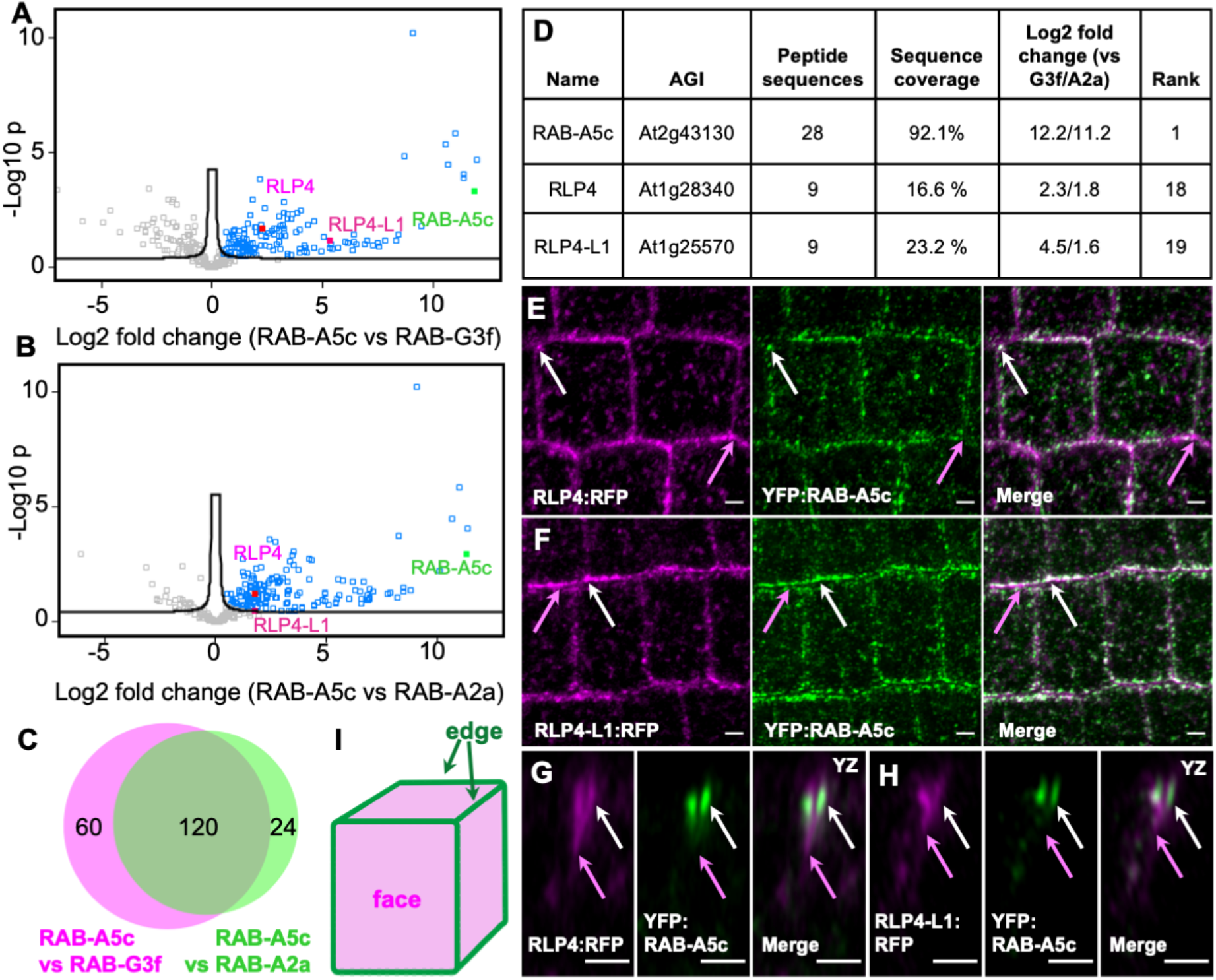
RLP4 and RLP4-L1 are interactors of RAB-A5c. **(A**,**B)** Volcano plots of the interactomes of YFP:RAB-A5c compared to YFP:RAB-G3f (A) and YFP:RAB-A2a (B). Proteins significantly enriched in the YFP:RAB-A5c interactome are colour-coded in blue. **(C)** Venn diagram of proteins enriched in the YFP:RAB-A5c proteome shown in (A,B). **(D)** Final ranking for RLP4 and RLP4-L1 from the comparative proteomics approach. **(E-H)** Confocal laser scanning microscopy (CLSM) maximum intensity projections (E,F) or YZ orthogonal projections (G,H) of Arabidopsis lateral root cells co-expressing *pUBQ10::RLP4s:RFP* (magenta) and *pRAB-A5c::YFP:RAB-A5c* (green).RLP4s:RFP co-localise with YFP:RAB-A5c at edge compartments (white arrows) and additionally label the cell periphery at edges (magenta arrows). **(I)** Cartoon depiction of a cell showing edges and faces. Scale bars 5 µm.

We expressed fluorescently tagged versions of RLP4 and RLP4-L1 (collectively referred to as RLP4s below) to examine their intracellular localisation pattern. When stably expressed under the control of the UBQ10 promoter, RLP4s:RFP co-localised with YFP:RAB-A5c at edge compartments (Figure 1E-H, white arrows) in young lateral roots of *Arabidopsis*. RLP4s:RFP additionally localised to the cell periphery adjacent to cell edge compartments (Figure 1E-H, pink arrows) and punctae distributed throughout in the cytoplasm. In orthogonal projections, it was apparent that both the peripheral signal was strongest adjacent to YFP:RAB-A5c-labelled cell edge compartments (Figure 1G-I). In dividing cells, RLP4s:RFP colocalised with YFP:RAB-A5c at the cell plate (Figure 3B, S3A,C).

The RLP4s localisation pattern was reproducible in N-terminally tagged secRFP:RLP4s expressed under the control of the UBQ10 promoter (*pUBQ10::secRFP:RLP4s*; Figure S1A,B), as well as in lines expressing RLP4s:GFP under their respective endogenous promoters (*pRLP4::RLP4:GFP* and *pRLP4-L1::RLP4-L1:GFP*, Figure S1C,D). At the tissue level, *pRLP4::RLP4:GFP* and *pRLP4-L1::RLP4-L1:GFP* were highly expressed in growing tissues of the root and shoot, an expression pattern matching that of *pRAB-A5c::YFP:RAB-A5c* (Figure S1E-J).

### RLP4 and RLP4-L are edge-polarised within the plasma membrane

To characterise the subcellular localisation of RLP4s, we quantitatively analysed the colocalization of RLP4s:RFP with a series of membrane markers including the Golgi marker ST:YFP (Boevink et al., 1998), the TGN/EE marker VHA-a1:GFP (Dettmer et al., 2006), YFP:RAB-A5c (Kirchhelle et al., 2016), and the plasma membrane marker YFP:NPSN12 (Geldner et al., 2009). We used a bespoke image analysis strategy that involved hysteresis-based filtering to eliminate cytosolic background signal, and quantified Manders’ colocalization coefficients (Manders et al., 1993) to evaluate the fractions of each fluorescent marker overlapping with the co-expressed RLP4s:RFP. This analysis revealed minimal colocalization of RLP4s:RFP with ST:YFP-labelled Golgi (Figure 2A,E, S2A; Manders’ colocalization coefficient 0.09±0.05 and 0.09±0.04 for RLP4:RFP and RLP4-L1:RFP, respectively), while the majority of the RLP4s:RFP signal was distributed in similar proportions between the VHA-a1:GFP-labelled TGN/EE (Figure 2B,F, S2B; Manders’ colocalization coefficient 0.23±0.06 and 0.26±0.09, respectively), YFP:RAB-A5c-labelled edge compartments (Figure 2C,G, S2C; Manders’ colocalization coefficient 0.23±0.08 and 0.29±0.10, respectively), and the YFP:NPSN12-labelled PM (Figure 2D,H, S2D; Manders’ colocalization coefficient 0.29±0.09 and 0.26±0.09, respectively).

**Figure 2:**
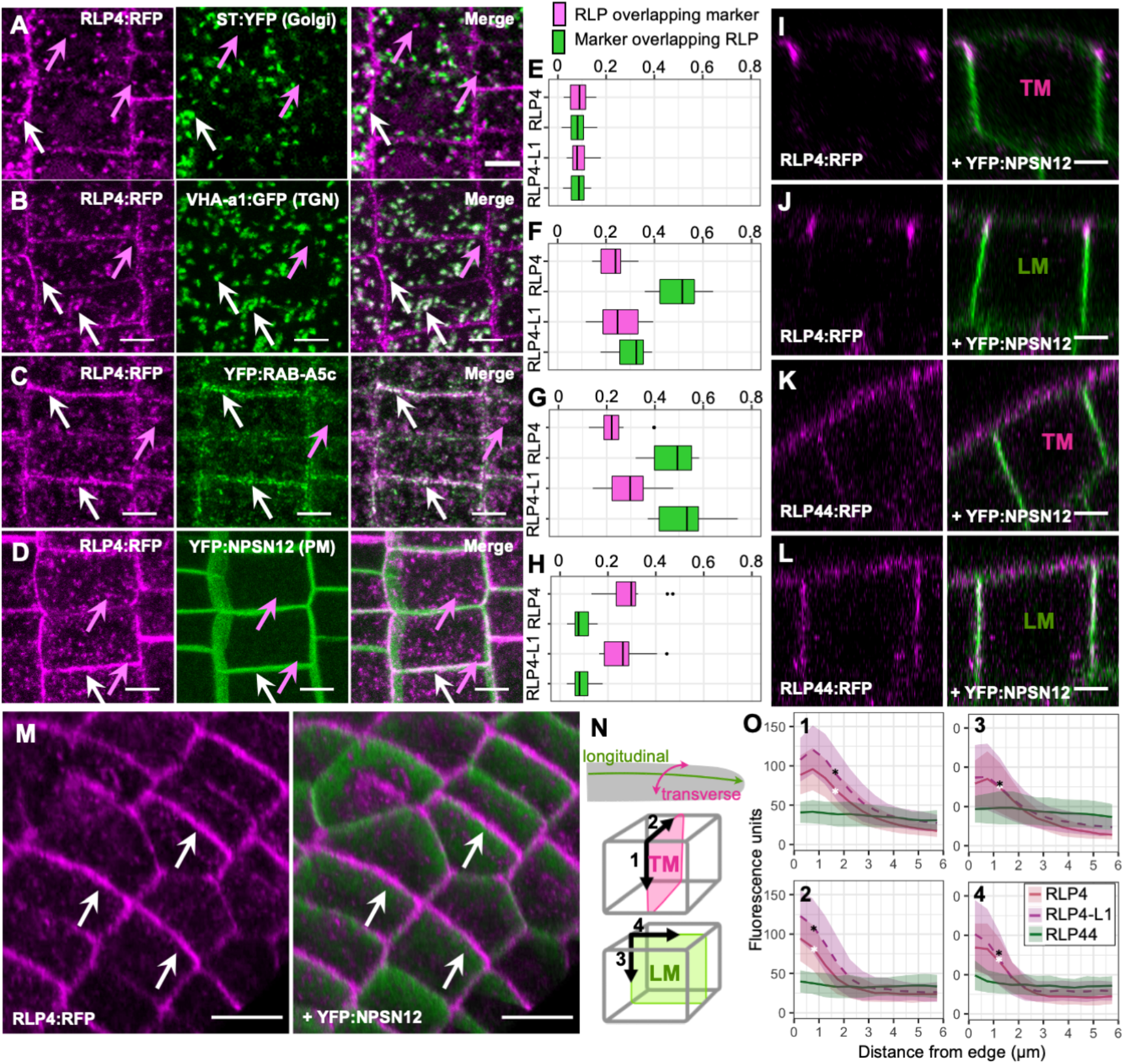
RLP4s are edge-polarised at the plasma membrane. **(A-D)** CLSM maximum intensity projections of lateral roots co-expressing pRLP4::RLP4:RFP and ST:YFP (A), VHA-a1:GFP (B), YFP:RAB-A5c (C), and pUBQ10::RLP4:RFP and YFP:NPSN12 (D). **(E-H)** Manders’ colocalization coefficients showing the fraction of RLP4:RFP and RLP4-L1:RFP colocalising with the membrane markers shown in (A-D). N≥13 regions from ≥4 roots. **(I-L)** CLSM XZ/YZ projections representing transverse (TM; I,K) and longitudinal (LM; J,L) midplane sections through meristematic lateral root cells co-expressing *pUBQ10::RLP4:RFP* (I,J) or *pUBQ10:RLP44:RFP* (K,L) and YFP:NPSN12. **(M)** Snapshot from MorphoGraphX (Barbier de Reuille et al., 2015) of a lateral root co-expressing RLP4:RFP and YFP:NPSN12. Note RLP4 is enriched at cell edges (white arrows). **(N)** Cartoon depiction of transverse (TM) and longitudinal (LM) midplane sections as those shown in (I-L). Black arrows depict trajectories along which fluorescence intensity was quantified as shown in (O): (1) longitudinal anticlinal, (2) longitudinal periclinal, (3) transverse anticlinal, (4) transverse periclinal. **(O)** Quantification of fluorescence intensity of RLP4:RLP (red), RLP4-L1:RFP (magenta, dashed line), and RLP44:RLP with increasing distance from the cell edge along the trajectories shown in (N). Lines indicate average fluorescence intensity (n ≥ 82 edges from 3 roots), shaded areas are +/-1SD. Note that RLP4:RFP and RLP4-L1:RFP were enriched at cell edges and fluorescence intensity rapidly decreased with increasing distance from the edge, whereas RLP44:RFP was uniformly distributed across the plasma membrane. Asterisks indicate distance from the cell edge at which RLP4:RFP (white asterisk) or RLP4-L1:RFP (black asterisk) signal intensity becomes significantly lower than at the edge (one-way ANOVA and post-hoc Tukey test, p<0.05). Scale bars 5 µm (A-D, I-L) or 10 µm (M).

Although a substantial fraction of RLP4s:RLP colocalised with YFP:NPSN12, only a small fraction of YFP:NPSN12 was labelled by RLP4s:RFP (Manders’ colocalization coefficient 0.09±0.04 in both cases). This further supported our qualitative observation that RLP4s:RFP localised to a subdomain of the PM at cell edges, which was particularly apparent in orthogonal or 3D projections of confocal stacks (Figure 2I,J,M, S1G,H, S2E-H). To quantitatively analyse the PM distribution of RLPs4:RFP, we used transverse (TM) and longitudinal (LM) midplane sections through meristematic root cells (Figure 2I,J,N, S2G,H). We quantified fluorescence intensity with increasing distance from cell edges in anticlinal and periclinal direction of RLP4s:RFP and RLP44:RFP (Figure 2K,L; supplementary table 2). RLP44 is a RLP that localises to the PM (Wolf et al., 2014) and is related to RLP4s, but does not localise to cell edge compartments. In our analysis, RLP44:RFP fluorescence was distributed relatively uniformly across cell faces (Figure 2O, green), whereas RLP4s:RFP fluorescence was brightest at cell edges and significantly decreased with increasing distance from cell edges in both anticlinal and periclinal direction (Figure 2O, red and magenta, respectively; asterisks indicate distance from edge from which point onwards signal intensity is significantly lower than at the edge itself (one-way ANOVA and post-hoc Tukey test, p<0.05)). Fluorescence intensity for RLP4s:RFP at edges was slightly higher in transverse sections than in longitudinal sections (92±29 vs 83±30 fluorescent units for RLP4:RFP and 117±38 vs 95±43 fluorescent units for RLP4-L1:RFP, respectively), indicating enrichment at longitudinal edges similar to the pattern previously described for YFP:RAB-A5c (Kirchhelle et al., 2019).

### Establishment of RLP4s cell edge polarity requires RAB-A5c activity

We next aimed to test how the polar RLP4s edge domain at the PM was established, and how different domains of the RLP4s protein may contribute to its localisation. RLP4s are predicted to contain a short intracellular domain, a transmembrane domain, and an extracellular domain carrying LRRs as well MLDs like those found in CrRLK1Ls ((Fritz-Laylin et al., 2005; Wang et al., 2008); Figure 3A). The extracellular domains of other RLPs have been shown to interact with extracellular proteinaceous ligands or the cell wall (Fiers et al., 2005; Wolf, 2022; Wolf et al., 2014), whereas the intracellular domain is expected to interact with intracellular trafficking machinery.

**Figure 3:**
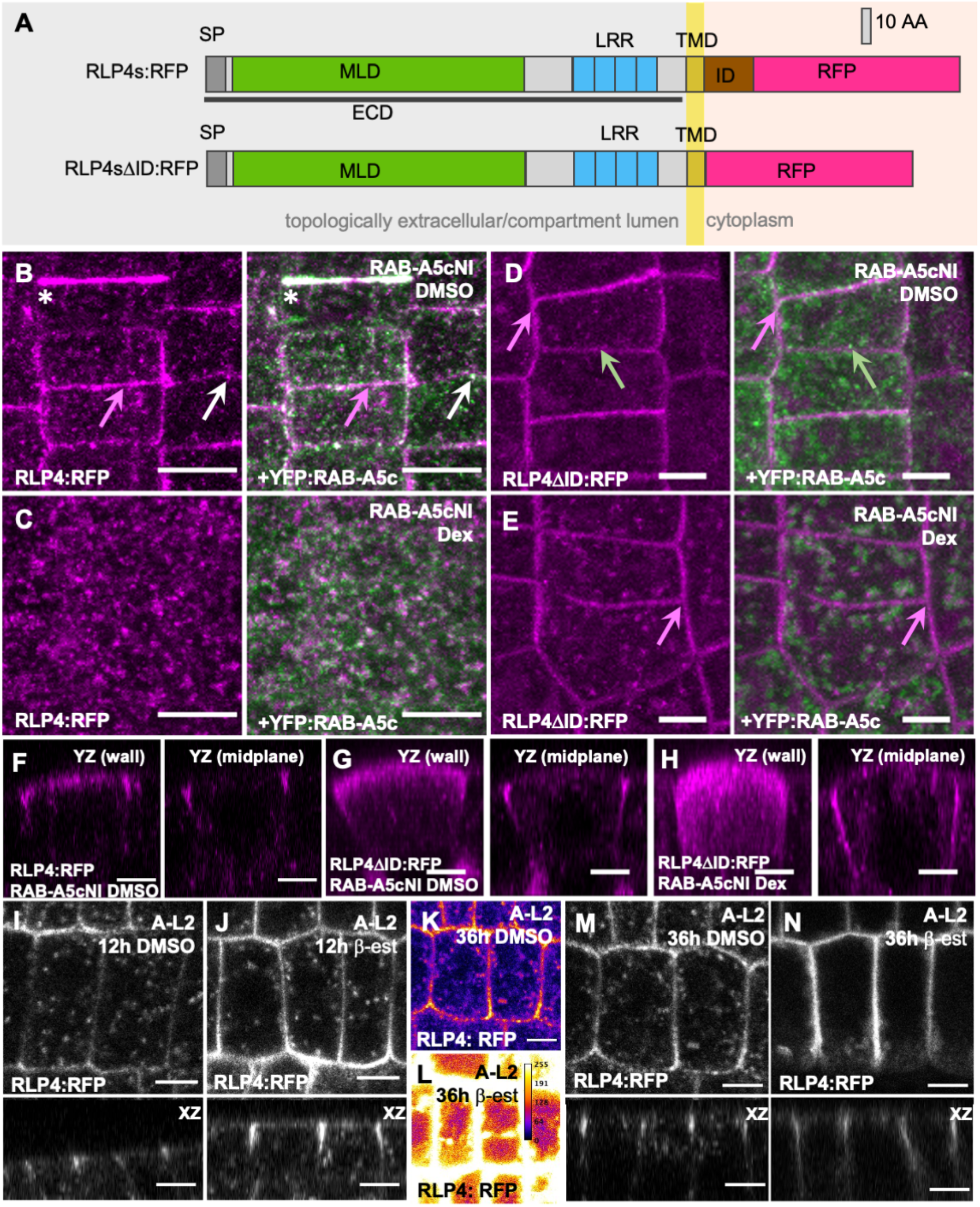
RLP4/RLP4-L1 cell edge polarity depends on cell wall association and RAB-A5c activity. **(A)** Cartoon depiction of RLP4s and their truncated variants used in this figure. **(B**,**C)** CLSM maximum intensity projections of lateral root cells co-expressing *pUBQ10::RLP4:RFP* (magenta), YFP:RAB-A5c (green), and Dex-inducible dominant-negative *AtRPS5a>>DEX>>RAB-A5c[N125I]* after 3d on DMSO (B) or 10µM Dex (C). Under control conditions (B), RLP4:RLP co-localised with YFP:RAB-A5c at the cell plate (asterisk) and cell edge compartments (white arrows), and additionally labelled the plasma membrane (magenta arrows). When RAB-A5c[N125I] was induced (C), RLP4:RFP and YFP:RAB-A5c localisation at cell edges was disrupted and shifted towards punctae distributed throughout the cytosol. **(D**,**E)** CLSM maximum intensity projections of lateral root cells co-expressing *pUBQ10::RLP4ΔID:RFP* (magenta), YFP:RAB-A5c (green), and Dex-inducible dominant-negative *AtRPS5a>>DEX>>RAB-A5c[N125I]* after 3d on DMSO (D) or 10µM Dex (E). Under control conditions (D), RLP4:RLP localised to the plasma membrane (magenta arrows), but did not co-localise with YFP:RAB-A5c at cell edge compartments (green arrows), When RAB-A5c[N125I] was induced (E), RLP4:RFP remained at the plasma membrane (magenta arrows). **(F-G)** CLSM YZ orthogonal projections of images as those shown in (B,D,E) showing RLP4:RFP localization (F) or RLP4*Δ*ID:RFP localisation (G,H) at transverse walls (left) or mid-plane sections without (F,G) or with (H) the induction of RAB-A5cNI. Note RLP4*Δ*ID:RFP is edge-polarised in the absence of RAB-A5cNI expression (G), but spreads out across cell faces when RAB-A5cNI is induced (H). **(J-M)** CLSM sections or XZ projections of lateral roots co-expressing *pUBQ10::RLP4:RFP* and β-estradiol-inducible A-L2 after 12h treatment with DMSO (J,L; control) or 10µM β-estradiol (K,M). **(I-N)** CLSM sections (top) or XZ orthogonal projections(bottom) of lateral roots co-expressing *pUBQ10::RLP4: RFP* and β-estradiol-inducible A-L2 after 12h (I,J) or 36h (K-N) treatment with DMSO (I,K,M) or 10µM β-estradiol (J,L,N). K and L are the same image displayed in gray-scale (M) or as a heat map (K) to emphasise intensity. Note after 36h A-L2 induction, intensity of RLP4:RFP at the cell surface was very high (compare L and K), precluding meaningful quantitative comparisons between induced and non-induced conditions with regard to localisation, so imagine parameters were adjusted to acquire images without saturation (N). Scale bars 5 µm.

YFP:RAB-A5c-labelled cell edge compartments were previously shown to not accumulate the endocytic tracer FM4-64 (Kirchhelle et al., 2016), suggesting that RAB-A5c is involved in *de novo* secretion, and RLP4s may be a cargo of this edge-directed secretory trafficking pathway. To test whether RAB-A5c activity was required to transport RLP4s:RFP to the cell surface, we inducibly expressed the dominant-negative protein variant RAB-A5c[N125I] to disrupt RAB-A5c function (*AtRPS5a>>DEX>>RAB-A5c[N125I]*, (Kirchhelle et al., 2016)). RAB-A5c[N125I] iwa predicted to competitively inhibit RAB-A5c activation through titration of essential regulatory interactors and was previously shown to functionally inhibit RAB-A5c, but did not disrupt bulk secretory trafficking to the cell surface in *Arabidopsis* primary and lateral roots (Kirchhelle et al., 2016).

We induced RAB-A5c[N125I] expression in plants co-expressing *pUBQ10::RLP4s:RFP* and *pRAB-A5c::YFP:RAB-A5c*, and examined lateral root meristematic cells after 3d induction (Figure 3B,C; Figure S3 A,B). In the presence of RAB-A5c[N125I], edge compartments labelled by RLP4s:RFP and YFP:RAB-A5c were no longer detectable, instead both markers labelled intracellular punctae distributed throughout the cytosol. Notably, RLP4s:RFP localisation at cell edges of the PM was almost completely abolished, suggesting that RLP4s:RFP localisation at the cell surface was dependent on RAB-A5c activity.

To test this observation further, we generated truncated versions of RLP4s:RFP that lacked their cytosol-facing intracellular domain, which is expected to interact with the cytosolic trafficking machinery including RAB-A5c (Figure 3A, *pUBQ10::RLP4sΔID:RFP*). These protein variants localised almost exclusively to the PM (Figure 3D, S3E), and did not substantially label cell edge compartments or other intracellular compartments. In contrast to full-length RLP4s:RFP, expression of *AtRPS5a>>DEX>>RAB-A5c[N125I]* did not abolish the localisation of RLP4sΔID:RFP to the PM (Figure 3E, S3F). This suggests RLP4sΔID:RFP reached the PM not through RAB-A5c-mediated trafficking, but through secretory bulk flow, and that the intracellular domain of RLP4s is required for secretion through edge-directed transport. Interestingly, under control conditions RLP4sΔID:RFP were enriched at cell edges in a pattern reminiscent of the full-length protein versions (Figure 3F,G, S3G). In the presence of RAB-A5c[N125I], RLP4sΔID:RFP localisation expanded from the cell edges towards the cell faces (Figure 3H, S3E), suggesting that edge-polarisation at the PM was dependent on RAB-A5c function even for protein variants that reached the cell surface independently of RAB-A5c-mediated transport.

It is well established that endocytosis and polar recycling are essential for the establishment of polarity domains at different cell faces of the PM (Geldner et al., 2001; Łangowski et al., 2016), so we also examined the effect of inhibiting endocytosis on RLP4s polarity at the PM. We used plants conditionally over-expressing the clathrin uncoating factor AUXILIN-LIKE2 (A-L2), which causes specific inhibition of clathrin-mediated endocytosis (Adamowski et al., 2018). After 12h induction of AL-2, RLP4s:RFP remained strongly enriched at cell edges (Figure 3I,J). When we extended the induction time to 36h, numerous PM invaginations formed and large quantities of RLP4s:RFP accumulated at PM (Figure 3K,L), necessitating an adjustment of imaging parameters to avoid saturation in order to examine the protein distribution. Even under these conditions, RLP4s:RFP were still visibly enriched at cell edges, albeit to a lesser extent than in controls (Figure 3M,N, Figure SI-J).

Taken together, our results show that RAB-A5c-mediated secretory transport is essential for the establishment of RLP4s cell edge polarity at the PM. Even RLP4s protein variants that reached the cell surface independently of RAB-A5c were polarized to cell edges only when RAB-A5c was active, suggesting these RLP4s variant may interact with other edge-specific factors deposited by RAB-A5c. In contrast to well-characterised examples of polarity at cell faces (e.g. PIN1), RLP4s polarity was not strictly dependent on endocytosis.

### RLP4s are stabilised at the cell surface through cell wall association

To functionally characterise the extracellular domain of RLP4s, we generated further protein truncations (Figure 4A). We generated RLP4s variants lacking their extracellular domains (ECD) fused to a secreted version of RFP (secRFP) to target them to the secretory pathway, as the truncation eliminated their intrinsic N-terminal signal peptide (*pUBQ10::secRFP:RLP4sΔECD*). The equivalently N-terminally tagged full-length proteins (*pUBQ10::secRFP:RLP4s)* localised in the same pattern as C-terminally tagged RLP4s (Figure S1C,D), suggesting that the N-terminal position of the tag did not interfere with protein transport. We expected that secRFP:RLP4sΔECD would still be able to interact with the intracellular trafficking machinery involved in transport of the full-length protein – however, secRFP:RLP4sΔECD exclusively localised to intracellular compartments in lateral roots, and did not label edge compartments or the PM (Figure 4B-D; Figure S3A-C). We quantitatively analysed the colocalization between secRFP:RLP4sΔECD and ST:YFP (Figure 4C,E, Manders’ colocalization coefficient 0.36±0.12 and 0.28±0.14, respectively) and VHA-a1:GFP (Figure 4D,F, Manders’ colocalization coefficient 0.44±0.17 and 0.51±0.11, respectively), and found colocalization with these markers was significantly increased in comparison to full-length RLP4s:RFP (Figure 4E,F vs Figure 2E,F; p<0.001, ANOVA and post-hoc Tukey test). By contrast, an equivalent truncation of the related RLP44, secRFP:RLP44ΔECD, labelled the PM rather than intracellular compartments (Figure S4D).

**Figure 4:**
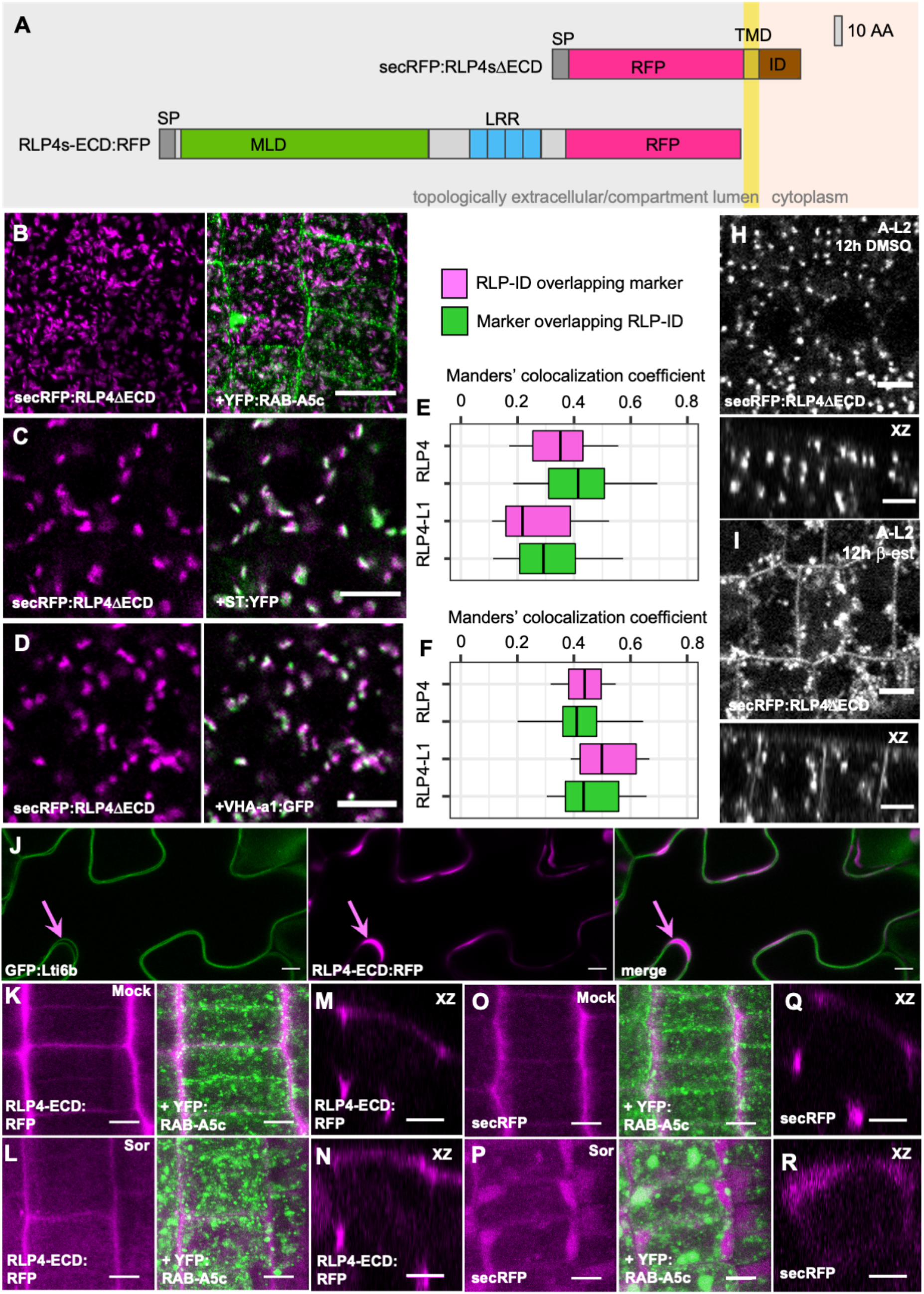
RLP4s are stabilised at the cell surface via cell wall association. **(A)** Cartoon depiction of truncated RLP4s variants used in this figure. **(B)** CLSM maximum intensity projections of lateral roots co-expressing *pUBQ10::RLP4ΔECD* and YFP:RAB-A5c. **(C**,**D)** CLSM maximum intensity projections of lateral roots co-expressing *pUBQ10::RLP4ΔECD* and ST-YFP (E), and VHA-a1:GFP (F). **(E**,**F)** Manders’ colocalization coefficients showing the fraction of RLP4:RFP and RLP4-L1:RFP colocalising with the membrane markers shown in (B,C). n ≥ 14 regions from ≥3 roots. **(F-I)** CLSM sections (top) or XZ orthogonal projections (bottom) of lateral roots co-expressing *pUBQ10::RLP4ΔECD* and β-estradiol-inducible A-L2 after 12h treatment with DMSO (H) or 10µM β-estradiol (I). **(J)** CLSM single optical section of leaf epidermal pavement cells co-expressing *p35S::GFP:Lti6b* and *pUB10:RLP4-ECD:RFP*. Magenta arrow indicates RLP4-ECD:RFP localisation in the apoplast between plasma membranes of two cells labelled by GFP:Lti6b. **(K-R)** CLSM maximum intensity projections (K,L,O,P) or XZ orthogonal projections (M,N,Q,R) of lateral roots co-expressing *pUBQ10::RLP4-ECD:RFP* (K-N) or *pUBQ10::secRFP* (O-R) and YFP:RAB-A5c after 30 minutes incubation in H_2_O (K,M,O,Q) or 500mM sorbitol (L,N,P,R). Scale bars 10 µm (B-D) or 5 µm (H-R).

We also treated seedlings expressing secRFP:RLP4sΔECD with the ARF-GEF inhibitor Brefeldin-A (BFA), which leads to the formation of so-called BFA bodies, large membrane aggregations that accumulate TGN/EE-localised proteins at their centre and Golgi-localised proteins at their periphery (Chow et al., 2008; Dettmer et al., 2006; Steinmann et al., 1999). As expected based on colocalization with TGN/EE and Golgi markers, secRFP:RLP4sΔECD labelled both the centre and periphery of BFA bodies (Figure S4E-H), but surprisingly, also faintly labelled the PM. It has been reported that inhibition of ARF-GEF function can lead to reduced internalisation of PM-localised proteins through inhibition of endocytosis (Irani et al., 2012; Naramoto et al., 2010; Teh and Moore, 2007). We therefore hypothesised that secRFP:RLP4sΔECD undergo constitutive secretion, but are rapidly endocytosed under standard growth conditions and therefore do not accumulate at detectable levels at the PM unless endocytosis is perturbed. To test this hypothesis, we co-expressed AUXILIN-LIKE1 or AUXILIN-LIKE2 (A-L1/A-L2) with secRFP:RLP4sΔECD to inhibit clathrin-mediated endocytosis (Adamowski et al., 2018). Induction of A-L1 and A-L2 expression resulted in partial relocalisation of secRFP:RLP4sΔECD to the PM (Figure 4H,I, Figure S4I-N), consistent with the hypothesis that secRFP:RLP4sΔECD undergo constitutive secretion and endocytosis. This suggests that the extracellular domain of RLP4s is required to maintain the proteins at the PM. In contrast to full-length RLP4s:RFP, secRFP:RLP4sΔECD were not edge-polarised at the PM in the presence of A-L2 (Figure 3J,N vs 4I), indicating the extracellular domain of RLP4s is required for edge polarisation.

The extracellular domain of RLP4s contains an MLD, a domain which is also found in RLKs of the CrRLK-L1 family, where it has been shown to bind pectins in vivo (Lin et al., 2022; Tang et al., 2022). To test whether RLP4s can also associate with the cell wall via their ECD, we expressed RLP4s variants that comprised solely of their ECD fused to RFP, while lacking their intracellular and transmembrane domains (*pUBQ10::RLP4s-ECD:RFP*).

These truncations were secreted to the apoplast, which was particularly obvious in leaf epidermal pavement cells, where cell walls are thick enough to resolve plasma membranes from neighbouring cells in confocal images (Figure 4J). In lateral roots, RLP4s-ECD:RFP were like-wise secreted, with the strongest signal emanating from cell edges (Figure 4K,M, S4O,P). This pattern was also observed for secRFP alone (Figure 4O,Q), and presumably reflects the geometry of the apoplast rather than specific targeting of the protein to cell edges. We next treated cells with 500 mM sorbitol for 30min to plasmolyse them. Under these conditions, secRFP flooded into the gap between the retracting protoplast and the cell wall (Figure 4P,R), whereas RLP4s-ECD:RFP remained at the cell wall (Figure 4L,N, S4Q,R), suggesting that the ECD of RLP4s can associate with the cell wall.

Taken together, this demonstrates that the ECD of RLP4s could associate with the cell wall to stabilise RLP4s at the PM. Our data also suggest the ECD of RLP4s is required for polarised localisation to cell edges, as truncations lacking this domain did not edge-polarise within the PM even when endocytosis was inhibited to allow accumulation at the PM (whereas the full-length protein was edge-localised under the same conditions, compare Figure 3J and 4I).

### RLP4s are required for directional growth control

We have previously shown that inhibition of RAB-A5c function disrupts directional growth in lateral root cells (Kirchhelle et al., 2016). To investigate the biological function of RLP4s and test whether RLP4s are required for edge-based growth control, we used CRISPR-Cas9 to obtain transcriptional null *rlp4, rlp4-l1* and *rlp4 rlp4-l1* mutants (Figure 5). Seedlings of each mutant line were phenotypically indistinguishable from wild-type Col-0 plants on standard growth conditions (Figure 5A,B).

**Figure 5:**
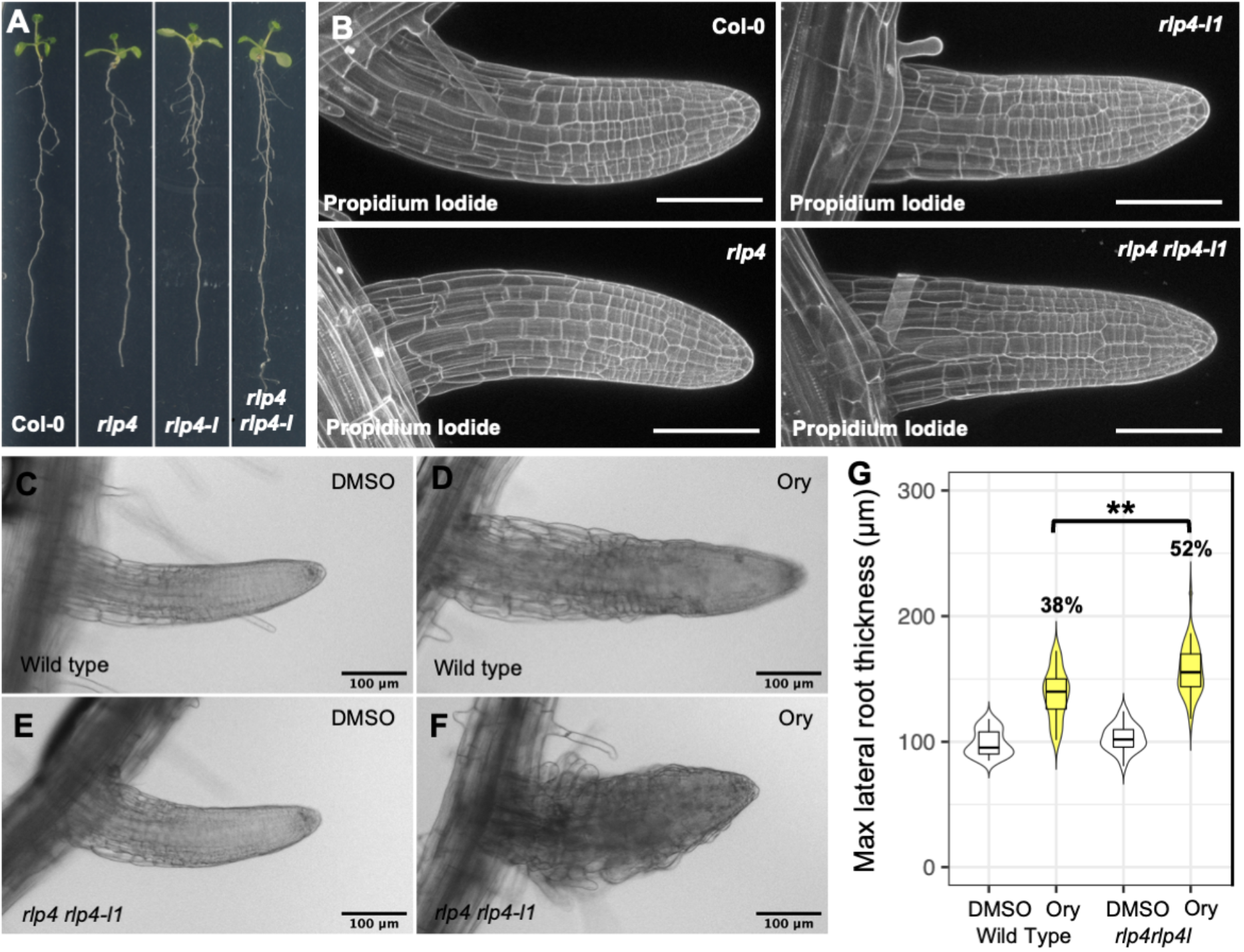
A *rlp4 rlp4-l1* mutant is hypersensitive to microtubule perturbation. **(A)** Photographs of 10d old Col-0, *rlp4, rlp4-l1*, and *rlp4 rlp4-l1* seedings. **(B)** CLSM maximum intensity projections of lateral roots from seedlings shown in (A). The cell wall was stained with propodium iodide. **(C-F)** Lateral roots from Col-0 Wild Type (C,D) and *rlp4 rlp4-l1* plants (E,F) 3 days after transfer to 0.1% DMSO (C,E) or 250 nM Ory (D,F). **(G)** Violin plots of the mean maximum diameter of lateral roots like those shown in (C-F) (n ≥ 35). Difference in diameter (%) between DMSO and 250nM Ory treatments for each genotype noted above respective columns. There is a significant difference in relative diameter increase (p=0.003) in response to Ory treatment in *rlp4 rlp4-l1* vs wild-type (t-test). Scale bars 100 µm.

We have shown previously that loss of edge-based growth control can partially be compensated through enhanced alignment of microtubules and consequently, cellulose, in lateral roots (Kirchhelle et al., 2019). As a consequence, plants in which edge-directed transport is inhibited are hypersensitive to pharmacological disruption of microtubule organisation with the microtubule-depolymerising drug oryzalin (Kirchhelle et al., 2019). To test whether a similar mechanism may be active in *rlp4 rlp4-l1*, we treated wild-type and *rlp4 rlp4-l1* mutant plants with oryzalin and quantified lateral root thickness as a proxy for directional growth, and found significantly increased swelling in *rlp4 rlp4-l1* compared to wild-type lateral roots (Figure 5C-G; 52% vs 38%, respectively), phenocopying the previously observed effect in the presence of RAB-A5c[N125I] (Figure 6N, S6C; (Kirchhelle et al., 2019)).

**Figure 6:**
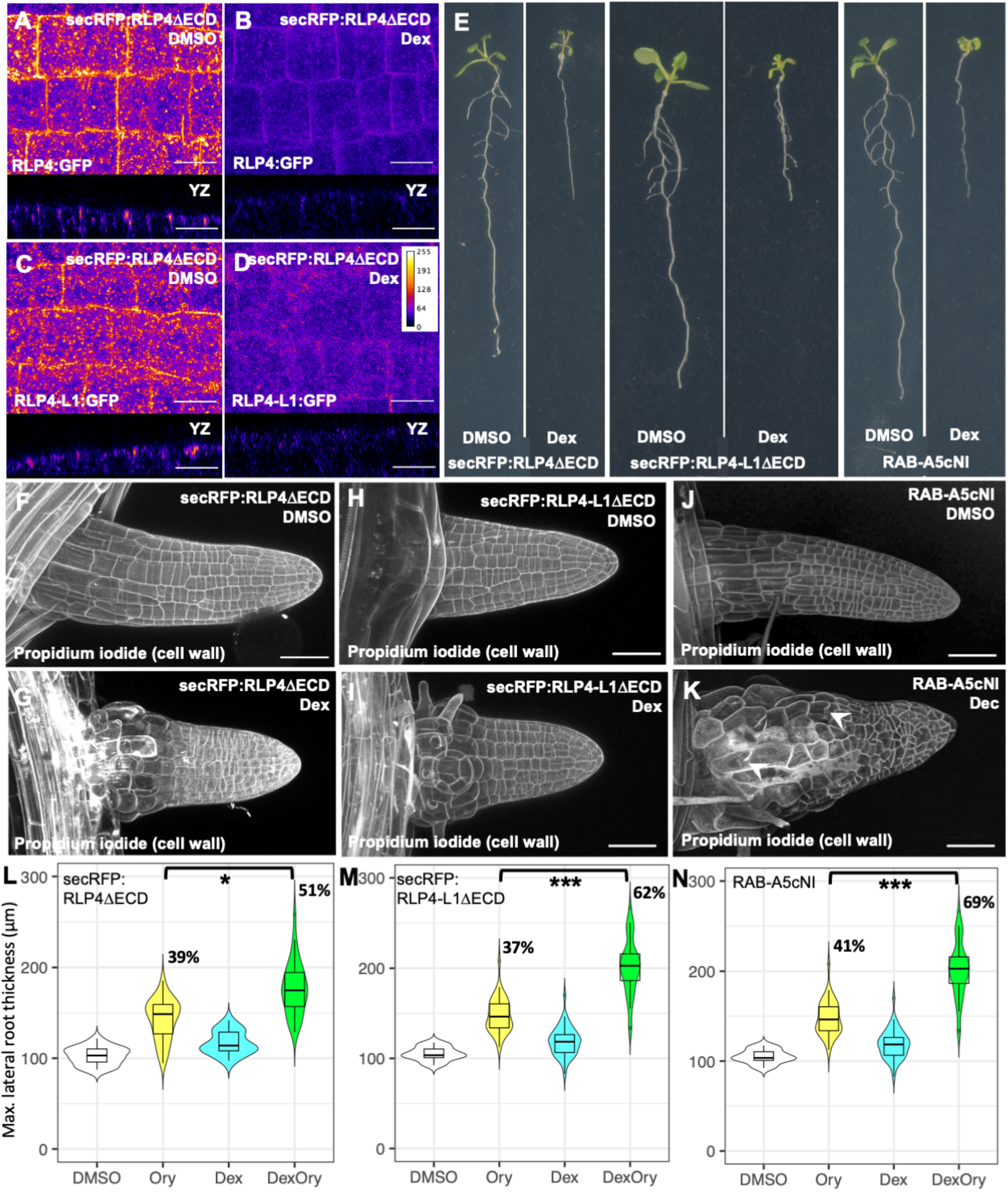
Inducible overexpression of RLP4ΔECD causes defects in directional growth, but not cell division. **(A-D)** CLSM maximum intensity heat map projections (top) and YZ orthogonal projections (bottom) of lateral roots from plants co-expressing *pRPS5a>>Dex>>RLP4ΔECD* and *pRLP4::RLP4:RFP* (A,B) or *pRLP4-L1::RLP4-L1:RFP* (C,D) 3 days after transfer to 0.1% DMSO (A,C) or 10 µM Dex (B,D). Note RLP4s:RFP intensity is decreased at cell edges and internal compartments after RLP4ΔECD induction. **(E)** Photographs of 10d old *pRPS5a>>Dex>>secRFP:RLP4ΔECD, pRPS5a>>Dex>>secRFP:RLP4-L1ΔECD* or *AtRPS5a>>DEX>>RAB-A5c[N125I]* seedlings grown on 0.1% DMSO (left) or 10 µM Dex (right). **(F-K)** CLSM maximum intensity projections of lateral roots from seedlings expressing *pRPS5a>>Dex>>secRFP:RLP4ΔECD* (F,G), *pRPS5a>>Dex>>secRFP:RLP4-L1ΔECD* (H,I) or *AtRPS5a>>DEX>>RAB-A5c[N125I]* (J,K) 3 days after transfer to 0.1% DMSO (F,H,J) or 10 µM Dex (G,I,K). The cell wall was stained with propidium iodide. **(L-N)** Violin plots of the mean maximum diameter of lateral roots from seedlings expressing *pRPS5a>>Dex>>secRFP:RLP4ΔECD* (L), *pRPS5a>>Dex>>secRFP:RLP4-L1ΔECD* (M) or *AtRPS5a>>DEX>>RAB-A5c[N125I]* (N). Lateral root diameter was measured after 3d on 0.1% DMSO (white), 250nM Oryzalin (yellow), 1 µM Dex (turquoise), or 250nM Oryzalin plus 1 µM Dex (green). n ≥ 18. There is a significant difference in relative diameter increase for all genotypes when treated with Ory in the presence of Dex compared to Ory treatment in the absence of Dex (*: p<0.05; ***: p<0.001, two-way ANOVA and post-hoc Tukey test). Scale bars 10 µm (A-D) or 10 µm (G-J).

We also aimed to conditionally disrupt RLP4s function, and hypothesised that overexpression of secRFP:RLP4sΔECD protein variants may disrupt transport of wild-type RLP4s through titration of the intracellular trafficking machinery. We expressed secRFP:RLP4sΔECD under the control of the Dex-inducible pOp/LhGR expression system (*AtRPS5a>>DEX>>secRFP:RLP4sΔECD;* (Craft et al., 2005; Samalova et al., 2019)), and examined whether secRFP:RLP4sΔECD overexpression did perturb *pRLP4::RLP4:GFP* and *pRLP4-L1::RLP4-L1:GFP* localization at cell edges. We found that RLP4s:GFP fluorescence intensity was strongly reduced at cell edges as well as intracellular compartments in the presence of overexpressed secRFP:RLP4sΔECD (Figure 6A-D, S5). When induced from germination onwards, secRFP:RLP4sΔECD caused growth defects reminiscent of those found in the root and in the shoot of plants expressing RAB-A5c[N125I] in 13/29 and 17/27 transgenic lines, respectively (Figure 6E). To examine the effect on directional growth specifically in lateral roots, we transferred 7-day-old seedlings grown under non-inducing conditions to Dex for 3 days. Under these conditions, lateral root morphology was strongly perturbed in *AtRPS5a>>DEX>>RAB-A5c[N125I]* (Figure 6J,K; (Kirchhelle et al., 2016)) and *AtRPS5a>>DEX>>secRFP:RLP4sΔECD* lines (Figure 6F-I).

We have previously demonstrated that expression of RAB-A5c[N125I] causes two independent defects in lateral roots: loss of directional growth in interphase cells and incomplete cell divisions (Figure 6K, arrowheads), the latter of which may be due to a conserved function of Rab-A GTPases or their interactors during cytokinesis (Chow et al., 2008; Kirchhelle et al., 2016). By contrast, Dex-inducible expression of secRFP:RLP4sΔECD did not cause any appreciable defects in cytokinesis, suggesting RLP4s play a role during interphase growth, but not cytokinesis. We furthermore tested whether *AtRPS5a>>DEX>>secRFP:RLP4sΔECD* lines were hypersensitive to oryzalin, and found that lateral root thickness after oryzalin treatment was significantly increased in plants expressing secRFP:RLP4sΔECD (Figure 6L,M, Figure S6A,B), phenocopying *AtRPS5a>>DEX>>RAB-A5c[N125I]* and *rlp4 rlp4-l1* ((Kirchhelle et al., 2019); Figure 5G, 6N, S6C).

## Discussion

In this study, we identify and characterise two Receptor-like Proteins, RLP4 and RLP4-L1, as components of edge-based growth control in plants.

We have demonstrated here that RLP4s occupy a unique polarity domain in the PM of epidermal cells in *Arabidopsis* lateral roots: their cell edges. This expands our current understanding of polarity domains in root epidermal cells, which recognises polar proteins localising to different cell faces (inner, outer, apical, and basal) as well as root hair initiation sites (Łangowski et al., 2016; Nakamura and Grebe, 2018). How cells establish and maintain different PM polarity domains is a much-studied question in the field. It is well established that constitutive endocytosis and rapid polar recycling are essential for the establishment of polarity domains at different cell faces (Geldner et al., 2001; Kleine-Vehn et al., 2011; Łangowski et al., 2016). The cell wall is likewise important for the establishment and maintenance of PM polarity domains at cell faces, which has been attributed to its effect on lateral diffusion of proteins in the PM (Feraru et al., 2011; Łangowski et al., 2016). By contrast, the role of polar secretion in establishing facial polarity is more contentious, in part due to the difficulty in experimentally separating *de novo* secretion and polar recycling. Based on computational modelling, it has been proposed that polar secretion contributes to the establishment of facial polarity (Łangowski et al., 2016). However, experimental inhibition of endocytosis abolishes facial polarity (Kitakura et al., 2011; Men et al., 2008), demonstrating that polar secretion alone is insufficient to generate facial polarity. By contrast, we show here that RLP4s polarity depends largely on RAB-A5c-mediated secretory transport rather than endocytosis. We show that while RLP4s undergo endocytosis, inhibition of endocytosis did not abolish edge localisation of full-length RLPs, suggesting that endocytosis was not required for establishment or maintenance of RLP4s polarity *per se*, but instead, largely affected overall protein levels at the cell surface. We also show that the cell wall association via their ECDs was required for RLP4s polarisation. Intriguingly, protein variants containing the ECD that reached the PM independently of RAB-A5 were still capable of edge-localisation, but this pattern was lost when RAB-A5c function was inhibited. This suggests that the ECD of RLP4s can interact with an edge-localised component whose presence is dependent on RAB-A5c. We have not yet identified specific interactors of RLP4s at the cell surface, but it is notable that RLP4s contain a Malectin-Like Domain (MLD) in their ECD, which are found in several cell-surface receptors implicated in sensing of cell wall composition or integrity (Boisson-Dernier et al., 2009; Deslauriers and Larsen, 2010; Guo et al., 2009; Hematy et al., 2007), and can bind the cell wall directly or indirectly (Du et al., 2018; Dünser et al., 2019; Feng et al., 2018; Lin et al., 2022; Moussu et al., 2018; Tang et al., 2022;). Differences in cell wall composition between edges and faces have been described in various cell types (Elliott and Kirchhelle, 2020; Parker et al., 2001; Parr et al., 1996; Suslov et al., 2009), and we have previously proposed that RAB-A5c is involved in local modification of cell wall properties at cell edges of lateral root epidermal cells (Kirchhelle et al., 2019). An attractive hypothesis based on these observations (Figure 7) is that RLP4s associate with an edge-enriched cell wall ligand at cell edges (which itself may be dependent on RAB-A5c-mediated transport) and the loss of edge-localised RLP4sΔID:RFP in the presence of RAB-A5c[125I] is a consequence of losing cell wall patterning at edges. Alternatively, it is possible that RLP4s localisation is determined by interaction with a proteinaceous binding partner at cell edges that is also a cargo of RAB-A5c, or is linked to local geometry at cell edges, which is disrupted in the presence of RAB-A5c[N125I].

**Figure 7:**
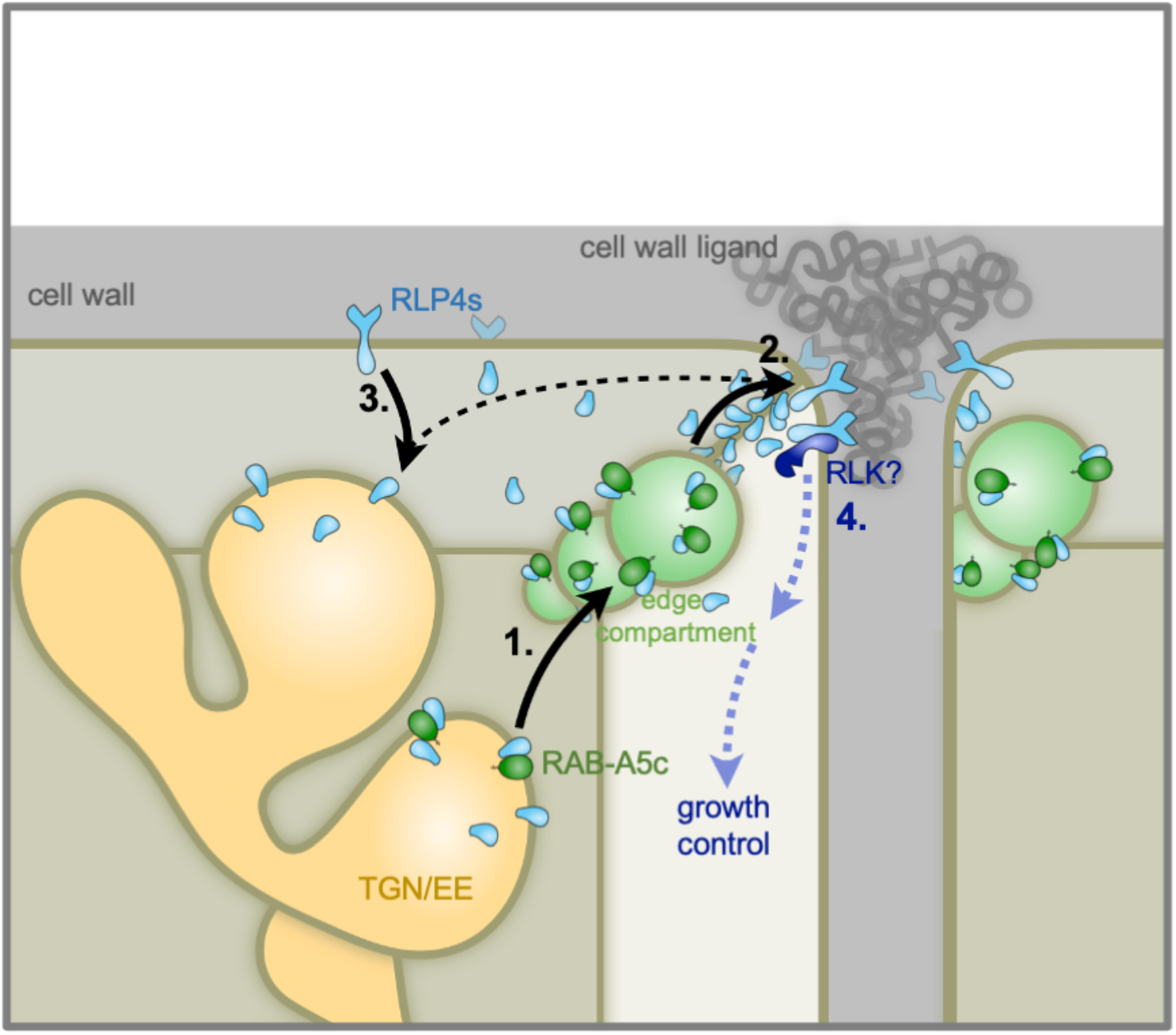
Model for RLP4s edge polarity establishment and function. 1.-3. Model for establishment of cell edge polarity. 1. RLPs (blue) are transported to from the TGN/EE to the PM at cell edges via cell edge compartments through association with RAB-A5c (green). 2. RLP4s are stabilised at cell edges through interaction with a cell wall ligand (grey). RLP4s that are not cell wall associated are rapidly endocytosed. **4**. Model for RLP4s function in growth control. RLP4s may act as a component of an edge-based signalling module through dimerization with an RLK.

In this study, we also demonstrate that RLP4s are functionally linked to control of directional growth. While a CRISPR knock-out mutant of RLP4s does not have a phenotype under standard growth conditions, it exhibits hypersensitivity to microtubule perturbation. The absence of directional growth defects in *rlp4 rlp4-l1* under standard growth conditions may be due to more efficient compensation of stable rather than induced loss of RLP4s at edges, but it is also possible that RAB-A5c contributes to directional growth control through mechanisms independent of RLP4s delivery to cell edges (for example, through local modification of the cell wall as we previously proposed). This raises the question of precisely how RLP4s contribute to growth control at cell edges? Answering this question conclusively is beyond the scope of this study but below, we offer a hypothesis based on our observations and previous findings regarding RLPs/RLKs in the literature.

Cell edges are a unique spatial domain from a mechanical perspective: they are exposed to high stresses for geometric reasons and in growing tissues, shear stresses may furthermore arise through differential growth rates in neighbouring cells (Jarvis et al., 2003; Kirchhelle et al., 2019; Roeder et al., 2022). This unique mechanical environment may explain the need to monitor the cell wall status specifically at cell edges through cell wall associated receptors like RLP4s. We therefore propose that RLP4s are components of an edge-localised signaling complex established in growing *Arabidopsis* tissues. Our data suggests that RLP4s are removed from the cell surface through endocytosis when they are not associated with the cell wall, suggesting that the subpopulation at the cell surface is proportional to its accessible cell wall ligand. This could provide a simple mechanism to directly monitor the biochemical composition of the cell wall at edges. It is also conceivable that RLP4s act as mechanosensors at edges: based on analogies to cell wall and extracellular matrix sensors in yeast and animal cells, it has been proposed that cell wall associated proteins like CrRLK1s may undergo molecular stretching in response to mechanical stress, which would in turn change their affinity for interacting proteins (Doblas et al., 2018). Like other RLPs, RLP4s lack an intracellular kinase domain to initiate a down-stream signaling cascade. However, it is well known that other RLPs interact with RLKs to form signaling modules at the cell surface and initiate intracellular signal cascades (Gust and Felix, 2014; Holzwart et al., 2018, 2020; Wolf et al., 2014). An RLP4s-based edge-localised signaling module containing an as-yet unidentified RLK(s) could thus monitor the biochemical or mechanical composition of cell edges during growth, linking the establishment and maintenance of spatio-temporal cell wall heterogeneity to polarised perception of the cell wall during plant growth.

Cells are 3D objects, and we often think of growth in terms of irreversible expansion of 3D volume. In growing tissues, growth is coordinated in different developmental zones, but often varies substantially in neighbouring cells (Hervieux et al., 2016; Kirchhelle et al., 2019; Uyttewaal et al., 2012). By contrast, growth at shared 2D faces must be strictly coordinated in neighbouring cells to maintain tissue integrity. Even cell faces that are not shared (i.e. at the outer organ surface) need to grow at appropriate rates to prevent cell bulging or rupture. 1D cell edges delimit cell faces in all directions, and requisite cell growth at faces can be considered as the integration of growth vectors along all edges delimiting the face. This implies that 2D and 3D growth patterns arise as a consequence of 1D growth control at cell edges (Korn, 1980). We and others have previously shown that cell edges are sites at which directional growth can be controlled (Ambrose et al., 2011; Kirchhelle et al., 2016), but here, we propose cell edges may also act as signalling domains at which positional information can be relayed and integrated into directional growth control. This provides a mechanistic framework to explain how tissue-level mechanics can be integrated into cellular growth through 1D cell edges. We have developed this framework of edge-based growth control in plant tissues, however, there are many conceptual parallels to epidermal tissues in animals. In such tissues, tricellular junctions (anticlinal edges) have been implicated in responses to mechanical stimuli, and also accumulate components of the JNK and Hippo growth signalling pathways (Bosveld et al., 2018). This opens up the intriguing possibility that 1D edges may function as sites of growth control in multicellular organisms of different lineages.

## Acknowledgements

This project was funded through Biotechnology and Biological Sciences Research Council (BBSRC) studentship 1810136 to LE, BBSRC grant BB/P01979X/1 to IM and CK, and Leverhulme Trust Early Career Fellowship ECF-2017-483 and ERC-2020-Stg 948514 — EDGE-CAM to CK. We thank Olivier Hamant and Yvon Jaillais for critical reading of this manuscript, and Mark Fricker for many fruitful discussions and advice.

## Author contributions

LE: Conceptualisation, Funding acquisition, Data curation, Investigation, Methodology, Resources, Writing – original draft, Writing – review and editing. MK: Investigation, Methodology. A-KS: Resources. Investigation. SW: Resources, Writing – review & editing. IM: Conceptualization, Funding acquisition, Project administration, Resources, Supervision. CK: Conceptualization, Data curation, Formal Analysis, Funding acquisition, Investigation, Methodology, Project administration, Supervision, Visualization, Writing – original draft, Writing – review & editing.

## Material and methods

### Plant materials and growth

The *Arabidopsis thaliana* ecotype Columbia was used throughout. We used various lines expressing fluorescent fusion proteins or dominant-negative proteins that have been described before: *pRAB-A5c::YFP:RAB-A5c* (Kirchhelle et al., 2016), *AtRPS5a>Dex>RAB-A5c[N125I]* (Kirchhelle et al., 2016) *pUBQ10::YFP:NPSN12* (Geldner et al., 2009), *pUBQ10::YFP:RAB-G3f* (Geldner et al., 2009), *pRAB:A2a::YFP:RAB-A2a* (Chow et al., 2008), *pVHA-a1::VHA-a1:GFP* (Dettmer et al., 2006), *p35S::ST:YFP* (Batoko et al., 2000) and *XVE>>AL1/XVE>>AL2* (Adamowski et al., 2018). For the simultaneous targeting of *RLP4* and *RLP4-L1* via CRISPR/Cas9, two suitable sequences for the generation of guide RNAs were determined using the ChopChop webpage (https://chopchop.cbu.uib.no/) and incorporated into oligonucleotides that also contained Eco31I recognition site at the 5’ end and pHEE2E-TRI-specific (Wang et al., 2015) sequence at the 3’ end. pHEE2E-TRI was used as template to amplify the two targeting sequences together with promoter and terminator regions. The amplified PCR product was gel purified and ligated into Eco31I (BsaI)-digested pHEE2E-TRI. The assembled construct was mobilised in *Agrobacterium tumefaciens* strain GV3101 and used to transform Col-0 plants. T1 plants were selected on ½ MS, 0.75 % phytoagar and 15 μg/mL hygromycin. Plates were covered with sheets of paper for four to six days until positive T1 plants with an elongated hypocotyl could be distinguished and kept for another four days at full light. Around 40 T1 plants were transferred to soil and analysed for mutations using primers. We isolated a Cas9-free double mutant with single base insertions in both genes (position 264 from ATG for RLP4, position 363 for RLP4-L1), leading to premature stop codons 14 and 11 exons downstream, respectively. All plants were grown at 20°C in a 16h:8h day:night cycle. Lateral roots were imaged 10 days after germination on upright half-strength Murashige and Skoog medium (MS, Sigma Aldrich) plates with 1% w/v sucrose and 0.8% Bacto agar (Appleton Woods) at pH 5.7. For conditional expression using either dexamethasone or β-estradiol, seedlings were grown for 7 days from germination before transfer to half-strength MS medium containing either 10 µM Dex (Sigma Aldrich – diluted from a 10 mM stock in DMSO), 10 µM β-estradiol (Sigma Aldrich – diluted from 10 mM a stock in DMSO) or an equivalent volume DMSO solvent for the indicated time period. Treatments with Brefeldin-A (BFA, Sigma Aldrich) were performed in half-strength MS liquid medium with 1% w/v sucrose at pH 5.7 using 50µM BFA (diluted from a 50mM stock in DMSO) for the indicated time periods. Plasmolysis was performed by immersion of plants in 0.5M sorbitol solution for 30 minutes. Introduction of novel transgenes into plants was performed using *Agrobacterium*-mediated floral dip transformation (Clough and Bent, 1998).

### Molecular Cloning

All genes were amplified by PCR using Phusion™ High-Fidelity DNA Polymerase (Thermo Fisher Scientific) from gDNA isolated from *Arabidopsis thaliana* ecotype Columbia-0. *pUBQ10::RLP4/4-L1:RFP, pUBQ10::RLP44:RFP, pUQ10B::RLP4/4-L1-ECD:RFP* and *pUB::RLP4/4-L1ΔID:RFP* were all generated by cloning the relevant gDNA region into pDONR207 (Invitrogen/Thermo Fisher Scientific) using Gateway™ BP Clonase II Enzyme Mix (Thermo Fisher Scientific) and subsequently into *pUB-RFP-DEST* (Grefen et al., 2010) using Gateway™ LR Clonase II Enzyme Mix (Thermo Fisher Scientific). For expression of RLP4:RFP and RLP4-L1:RFP from their native promoters, the UBQ10 promoter was removed from *pUB-RFP-DEST* through digestion with restriction endonucleases PspXI and PmeI (New England Biolabs) and the vector subsequently re-ligated using Klenow polymerase (DNA Polymerase I, Large fragment; New England Biolabs) and T4 DNA ligase (Thermo Fisher Scientific) to generate *pX-DEST-RFP*. The promoter region, 5’-UTR and coding region of RLP4 and RLP4-L were then amplified by PCR as single cassettes and cloned into *pDONR207* and eventually *pX-DEST-RFP* as described above. To generate *pUBQ10:secRFP:RLP4s* and *pUBQ10::secRFP:RLP4sΔECD*, the relevant gDNA regions were overlapped with *secRFP* (Samalova et al., 2006) and the cassettes cloned into *pENTR/D-TOPO* using a pENTR™/D-TOPO™ Cloning Kit (Thermo Fisher Scientific) and subsequently *pUB-DEST* (Grefen et al., 2010). For conditional expression of RLP4s and truncated variants using the pOp/LhGR system, transgenes were cloned into *pDONR207* using Gateway™ BP Clonase II Enzyme Mix (Thermo Fisher Scientific) and subsequently into *pOpIN2-RPS5a* (Samalova et al., 2019) using Gateway™ LR Clonase II Enzyme Mix (Thermo Fisher Scientific). All constructs were verified by Sanger sequencing (Source Bioscience) and restriction digests. For molecular cloning, *Escherichia coli* strains DH5α and DB3.1 were used. For *Agrobacterium*-mediated transformation of *Arabidopsis*, constructs were introduced into *Agrobacterium tumefaciens* strain GV3101::pMP90 by electroporation.

### Protein extraction and Proteomics

Co-immunoprecipitation and mass spectrometry for identification of interactors of YFP:RAB-A5c, YFP:RAB-A2a and YFP:RAB-G3f was performed as previously described (Kalde et al., 2019). In brief, co-IP experiments were carried out by isolating total microsomes from Arabidopsis roots expressing YFP:RAB-A5c (Kirchhelle et al., 2016), YFP:RAB-A2a (Chow et al., 2008) and YFP:RAB-G3f (Geldner et al., 2009), or no transgene (Col-0), as described in (Kalde et al., 2019). In-gel trypsin digest and mass spectrometry were performed by the Central Proteomic Facility, University of Oxford (www.proteomics.ox.ac.uk) and label-free quantification of the proteome was performed on three biological replicates using the SinQ pipeline (Trudgian et al., 2011). We excluded all proteins that did not occur in all three replicates of YFP:RAB-A5c, replaced all remaining zero values in the matrix by the half-minimum value across all detected proteins, and analysed the resulting 315 proteins for enrichment in RAB-A5c vs RAB-A2a and RAB-G3f proteomes using the Volcano plot function in the Perseus computational platform (Tyanova et al., 2016), with a S0 of 2 and FDR of 0.2, which identified 120 proteins significantly enriched in the YFP:RAB-A5c interactome compared to both YFP:RAB-A2a and YFP:RAB-G3f. We ranked these according to four different criteria: 1. Abundance in YFP:RAB-A5c interactome (descending order), 2. Relative enrichment against YFP:RAB-A2a interactome (descending order), 3. Relative enrichment against YFP:RAB-G3f interactome (descending order), and 4. Abundance in the Col-0 negative control (ascending order), and assigned a super rank according to the sum of individual ranks in ascending order (Supplementary table 1).

### Microscopy and image analysis

Confocal microscopy was performed using a Zeiss 880 CLSM using a C-Apochromat 40x/1.20 W Corr M27 objective or a Zeiss 980 CLSM using a C-Apochromat 40x/1.20 W Corr M27 objective. GFP, YFP, RFP, and PI were imaged as described before (Chow et al., 2008). Image analysis and processing (orthogonal sectioning, maximum intensity projections, image assembly, and quantification) was performed using Fiji (Schindelin et al., 2012). For the quantification of colocalization between RLP4s:RFP and various endomembrane markers, 3D regions with a length of 25µm in X/Y were selected from CLSM stacks of lateral roots. Background signal was removed using a hysteresis filter (Schindelin et al., 2012), using thresholds based on mean and minimum intensity -2SD of 10 randomly measured compartments for the respective CLSM channel, and Manders’ colocalization coefficients (Manders et al., 1993) were determined using the JACoP (Just Another Colocalisation Plugin) in Fiji (Bolte and Cordelieres, 2006). For the quantification of RLP4s:RFP at the PM, CLSM stacks of lateral roots co-expressing pUBQ10::YFP:NPSN12 and pUBQ10::RLP4s:RFP or pUBQ10::RLP44:RFP were collected at Nyquist resolution (voxel size 99.5nmx99.5nmx550nm). Midplane transverse and longitudinal sections of meristematic cells were generated in Fiji, and cellular outlines were manually traced using the PM marker YFP:NPSN12 as a reference. A plot profile with a width 7 pixels was generated and RFP intensity was measured along the profile. Average signal intensity for ≥82 edges from 3 roots was calculated for 0.5µm wide intervals starting at the edge for longitudinal anticlinal, transverse anticlinal, longitudinal periclinal and transverse periclinal walls, respectively. Average intensity +/-SD was plotted using the ggplot2 function in R (Wickham, 2009). To quantify root thickness, bright-field images of lateral roots were imported into Fiji, both sides of the root were traced manually along their longitudinal axis, and XY Cartesian coordinates for each pixel on the outline trace were exported as csv files and imported into RStudio (https://www.rstudio.com/). For each pixel on one side, the closest neighbour on the other side was determined and the Euclidian distance between pixels calculated using the nn2 function in the RANN package (https://CRAN.R-project.org/package=RANN). The maximum diameter of each root was calculated as the average of ten largest values excluding the tip-most 100 µm of each root to exclude the tapering tip.

### Statistical data analysis and plotting

Two-way ANOVA (analysis of variance) was performed in R using the aov function from the stats package (R Core Team, 2013). Tukey’s test was performed in R using the TukeyHSD function from the stats package, Student’s t-test were performed in R using the t.test function from the stats package. Box-, Ribbon-, and Violin-plots were generated in R using the ggplot2 function in R (Wickham, 2009).

