## supplementary figures for "Edge-based growth control in *Arabidopsis* involves two cell wall-associated Receptor-Like Proteins"

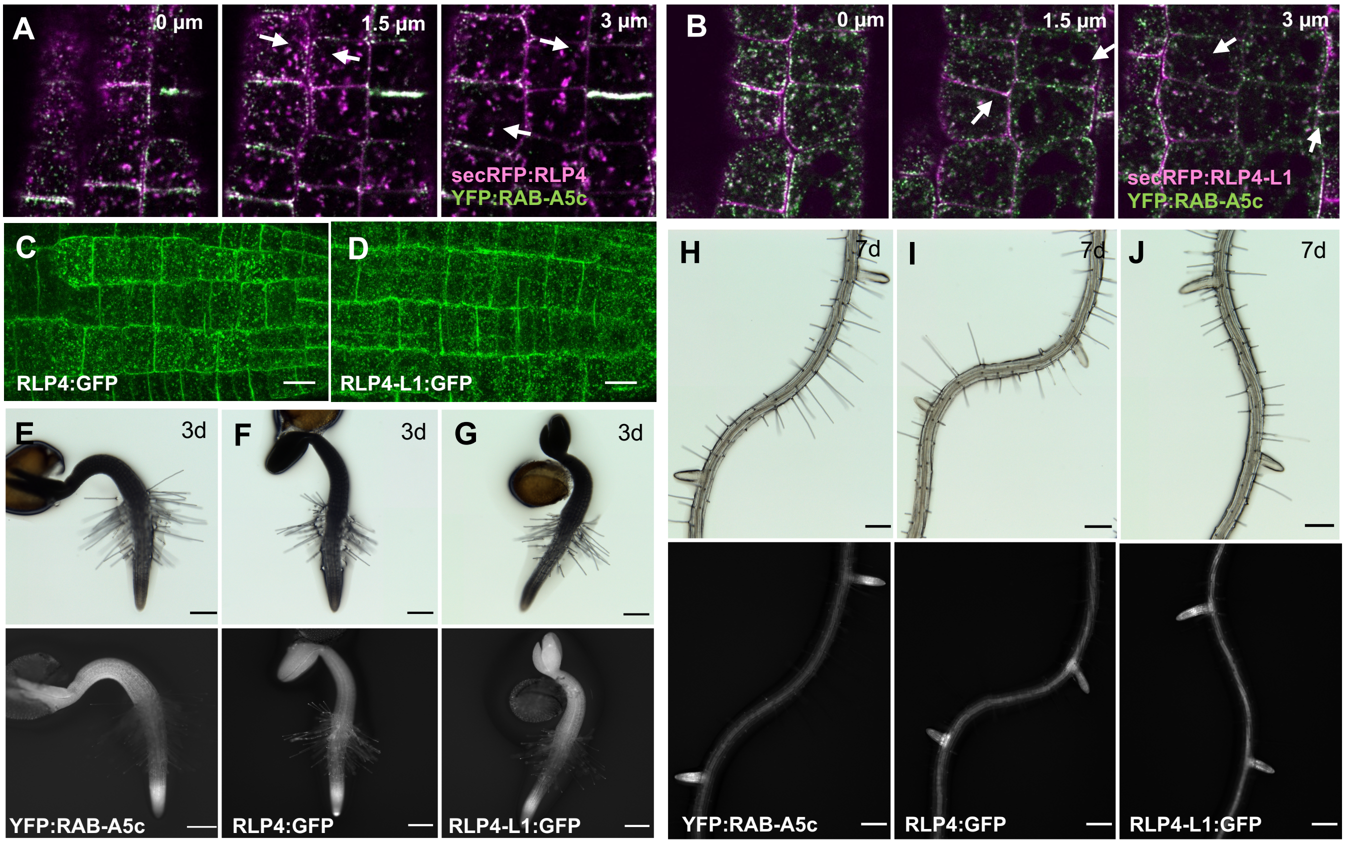


**Figure S1 (related to Figure 1): Localization and expression pattern of RLP4 and RLP4-L1**. **(A,B)** Sequential CLSM images of lateral root cells co-expressing *pUBQ10::secRFP:RLP4s:RFP* (magenta) and *pRAB-A5c::YFP:RAB-A5c* (green). **(C,D)** CLSM maximum intensity projections of lateral root cells expressing *pRLP4::RLP4:GFP* (C) or *pRLP4-L1::RLP4-L1:GFP* (D). **(E-J)** Brightfield and wide-field fluorescent images of 3 day old seedlings (E-G) or 10 day old roots (H-J) expressing *pRAB-A5c::YFP:RAB-A5c, pRLP4::RLP4:GFP* or *pRLP4-L1::RLP4-L1:GFP.* Scale bars 10µm (A-D), or 100µm (E-J).

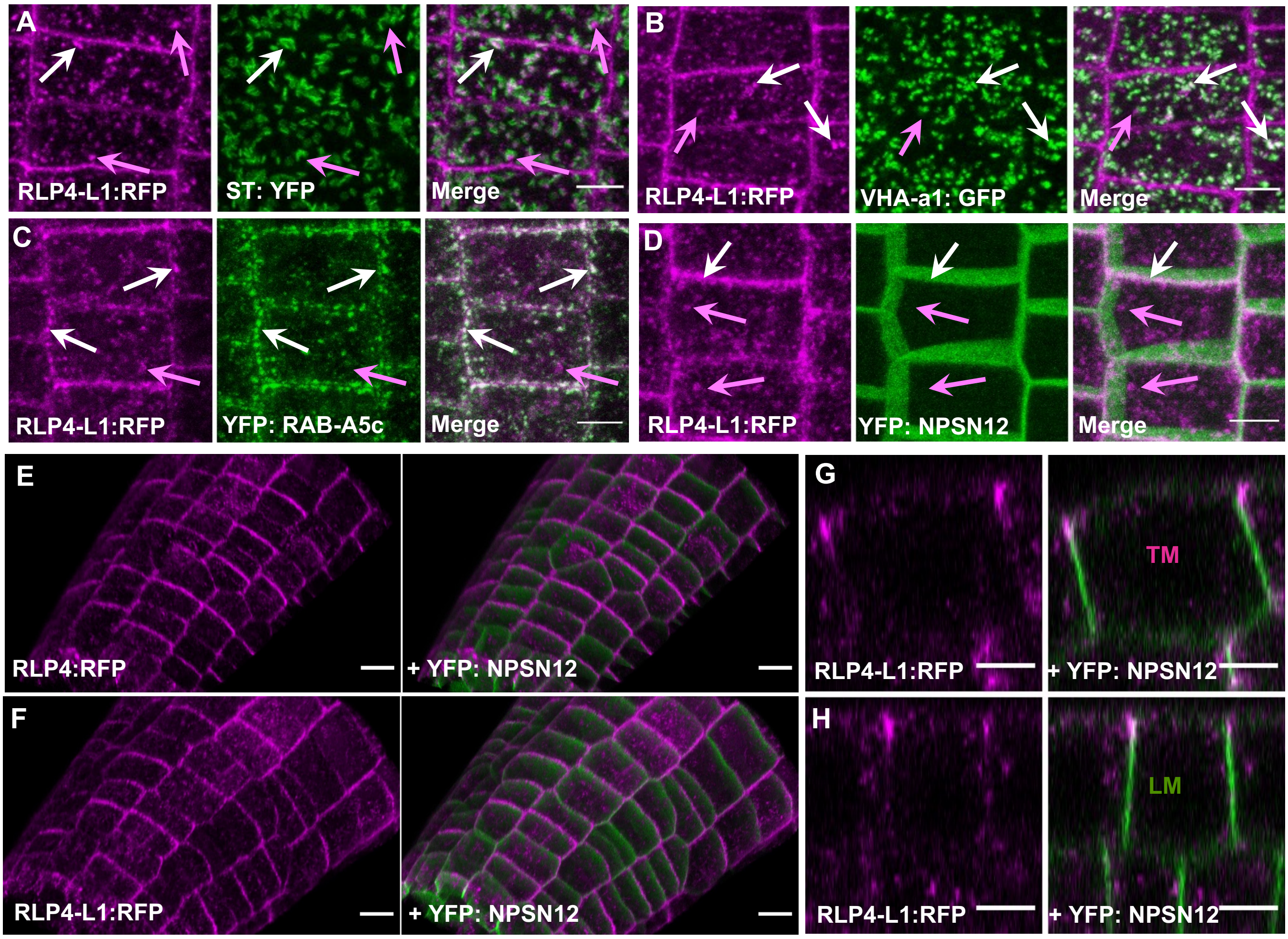


**Figure S2 (related to Figure 2): RLP4s are edge-restricted in *Arabidopsis* lateral root cells. (A-D)** CSLM projections of lateral root epidermal cells co-expressing *pRLP4-L1::RLP4-L1:RFP* and ST:YFP (A), VHA-a1:GFP (B), YFP:RAB-A5c (C), or *pUBQ10::RLP4-L1:RFP* with YFP:NPSN12 (D). **(E, F)** MorphoGraphX projections of lateral roots co-expressing *pUBQ10::RLP4s:RFP* (magenta) and YFP:NPSN12 (green). **(G,H)** CLSM XZ/YZ projections representing transverse (TM; G) and longitudinal (LM; H) midplane sections through meristematic lateral root cells co-expressing *pUBQ10::RLP4-L1:RFP* (magenta) and YFP:NPSN12 (green). Scale bars 5 µm (A-D,G,H) or 10 µm (E,F).

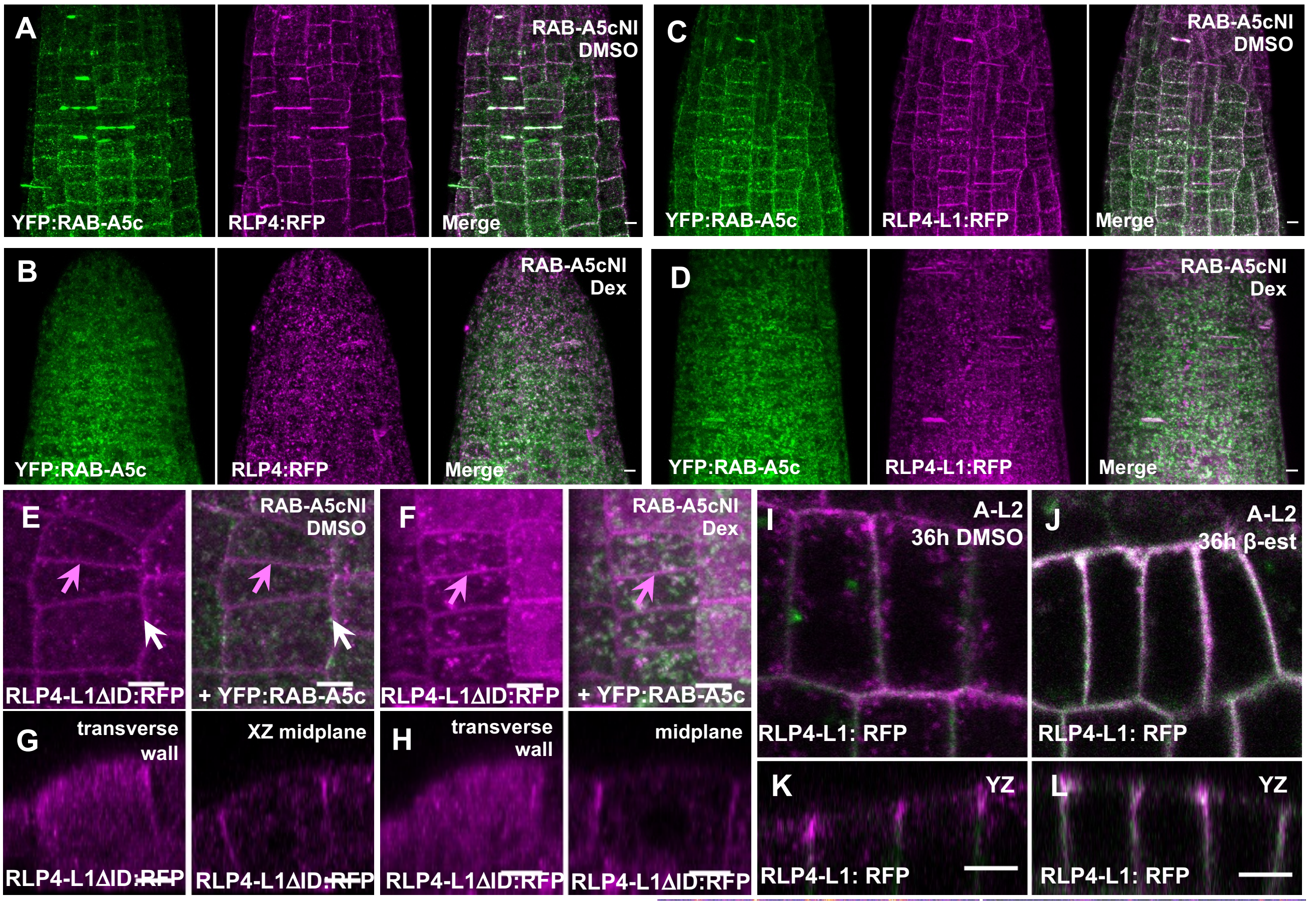


**Figure S3 (related to Figure 3): RLP4s edge polarity depends on RAB-A5c secretory transport, but not on endocytosis or rapid BFA-sensitive recycling. (A-D)** CLSM maximum intensity projections of lateral root cells co-expressing *pUBQ10::RLP4s1:RFP* (magenta), YFP:RAB-A5c (green), and Dex-inducible dominant-negative *AtRPS5a>>DEX>>RAB-A5c[N125I]* on DMSO (A,C) or 10μM Dex (B,D). Note that (A-D) are un-cropped versions of images shown in Figure 3B,C. **(E,F)** CLSM maximum intensity projections of lateral root cells co-expressing *pUBQ10::RLP4-L1ΔID:RFP* (magenta), YFP:RAB-A5c (green), and Dex-inducible dominant-negative *AtRPS5a>>DEX>>RAB-A5c[N125I]* after 3d on DMSO (E) or 10μM Dex (F). Under control conditions (D), RLP4:RLP localised to the plasma membrane (magenta arrows), but did not co-localise with YFP:RAB-A5c at cell edge compartments (white arrows), When RAB-A5c[N125I] was induced (F), RLP4:RFP remained at the plasma membrane (magenta arrows). **(G,H)** CLSM YZ orthogonal projections of images as those shown in (E,F) showing RLP4-L1ΔID:RFP localisation at transverse walls (left) or mid-plane sections without (G) or with (H) the induction of RAB-A5cNI. ). **(I-L)** CLSM sections (I,J) or XZ orthogonal projections (K,L) of lateral roots co-expressing *pUBQ10::RLP4-L1:RFP* (magenta), YFP:NPSN12 (green), and β-estradiol-inducible A-L2 after or 36h treatment with DMSO (I,) or 10μM β-estradiol (J,L). Note after 36h A-L2 induction, intensity of RLP4-L1:RFP at the cell surface was very high, precluding meaningful quantitative comparisons between induced and non-induced conditions with regard to localisation, so imagine parameters were adjusted to acquire images without saturation (J,L). Scale bars 5μm.

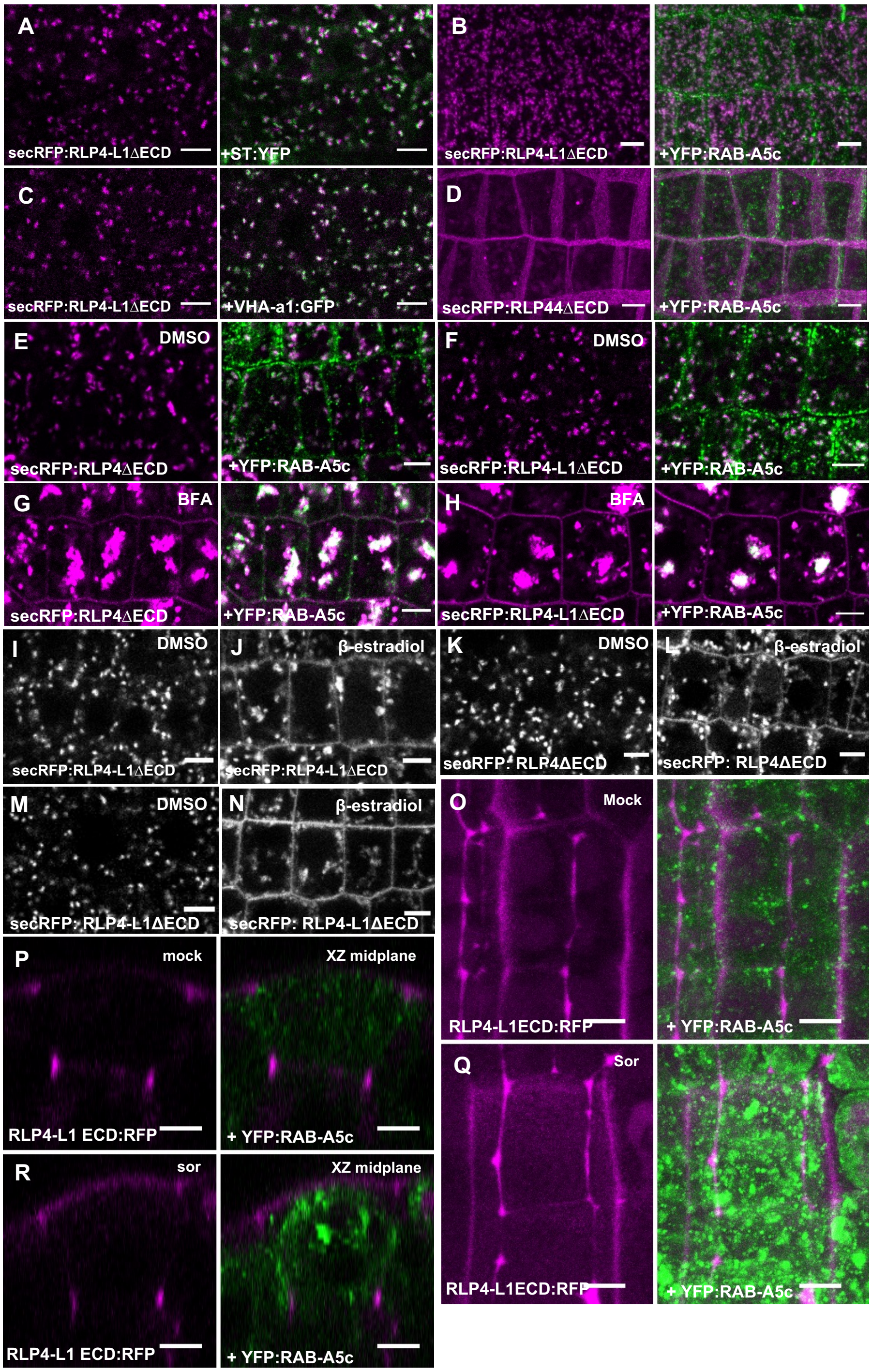


**Figure S4 (related to Figure 4): Cell wall association is required for RLP4s edge polarisation (A-C)** CLSM maximum intensity projections of lateral roots co-expressing *pUBQ10::RLP4-L1ΔECD* and ST-YFP (A), YFP:RAB-A5c (B), and VHA-a1:GFP (C). ). **(D)** CSLM maximum intensity projection of lateral root cells co-expressing *pUBQ10::secRFP:RLP44ΔECD* (magenta) and YFP:RAB-A5c (green). **(E-H)** CSLM images of lateral root cells co-expressing *pUBQ10::secRFP:RLP4ΔECD* (E,G; magenta) *or pUBQ10::secRFP:RLP4-L1ΔECD* (F,H; magenta) and YFP:RAB-A5c (green) after 30 minutes treatment with DMSO (E,F) or 50μM BFA (G,H). **(I-N)** CSLM images of lateral root cells co-expressing *pUBQ10::secRFP:RLP4-L1ΔECD* and ß-estradiol-inducible AUXILIN-LIKE1 (I,J) or ß-estradiol-inducible AUXILIN-LIKE2 (M,N), or *pUBQ10::secRFP:RLP4ΔECD* and ß-estradiol-inducible AUXILIN-LIKE1 (K,L) after 12 hours of treatment with DMSO (I,K,M) or 10μM ß-estradiol (J,L,N). **(O,P)** CLSM YZ projections of lateral root cells co-expressing *pUBQ10::RLP4-L1:RFP* and ß-estradiol-inducible AUXILIN-LIKE2 12 hours after transfer to DMSO (O) or 10μM ß-estradiol (P). **(O-R)** CSLM maximum intensity projections (O,Q) or XZ orthogonal projections (P,R) of lateral root cells co-expressing *pUBQ10::RLP4-L1ECD:RFP* (magenta) and YFP:RAB-A5c (green) after 30mins treatment with $H_2O$ (),P) or 0.5M sorbitol (Q,R). Scale bars 5μm.

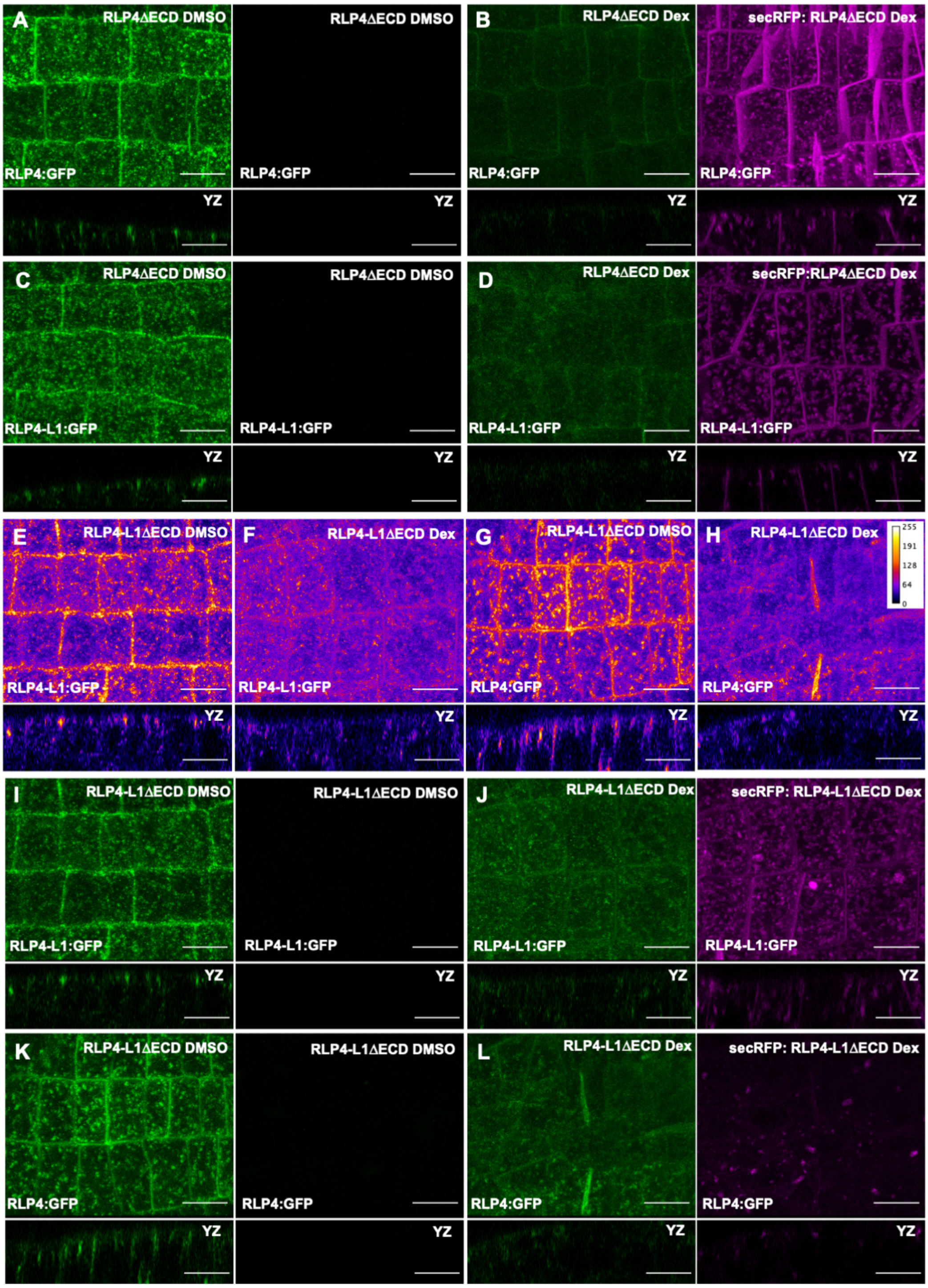


**Figure S5 (related to Figure 5): Inducible over-expression of RLP4/4-L1 truncated variants removes full-length RLP4/4-L1-GFP from cell edges. (A-D)** CSLM maximum intensity projections or YZ projections of lateral root cells co-expressing *(pRLP4::RLP4:GFP* (A,B; green) or *pRLP4-L1::RLP4-L1:GFP* (C,D; green) and *pUBQ10::secRFP:RLP4ΔECD* (magenta) 72 hours after transfer to DMSO (D,F) or 10μM Dex (E, G). Note images correspond to the images shown in Figure 5G-J using the ImageJ "fire" LUT. **(E-H)** CSLM maximum intensity projections or YZ projections of lateral root cells co-expressing *pRLP4::RLP4:GFP* (G,H) or *pRLP4-L1::RLP4-L1:GFP* (E,F) and *pUBQ10::secRFP-RLP4-L1ΔECD* 72 hours after transfer to DMSO (E,G) or 10μM Dex (G,H). Images are displayed in the ImageJ "fire" LUT to emphasize differences in intensity. **(I-L)** CSLM maximum intensity projections or YZ projections of lateral root cells co-expressing co-expressing *pRLP4::RLP4:GFP* (J,L; green) or *pRLP4-L1::RLP4-L1:GFP* (I,K; green) and *pUBQ10::secRFP-RLP4-L1ΔECD (magenta)* 72 hours after transfer to DMSO (L, N) or 10μM Dex (M, O). Note images correspond to the images shown in Figure S5E-H using the ImageJ "fire" LUT. Scale bars 10μm.

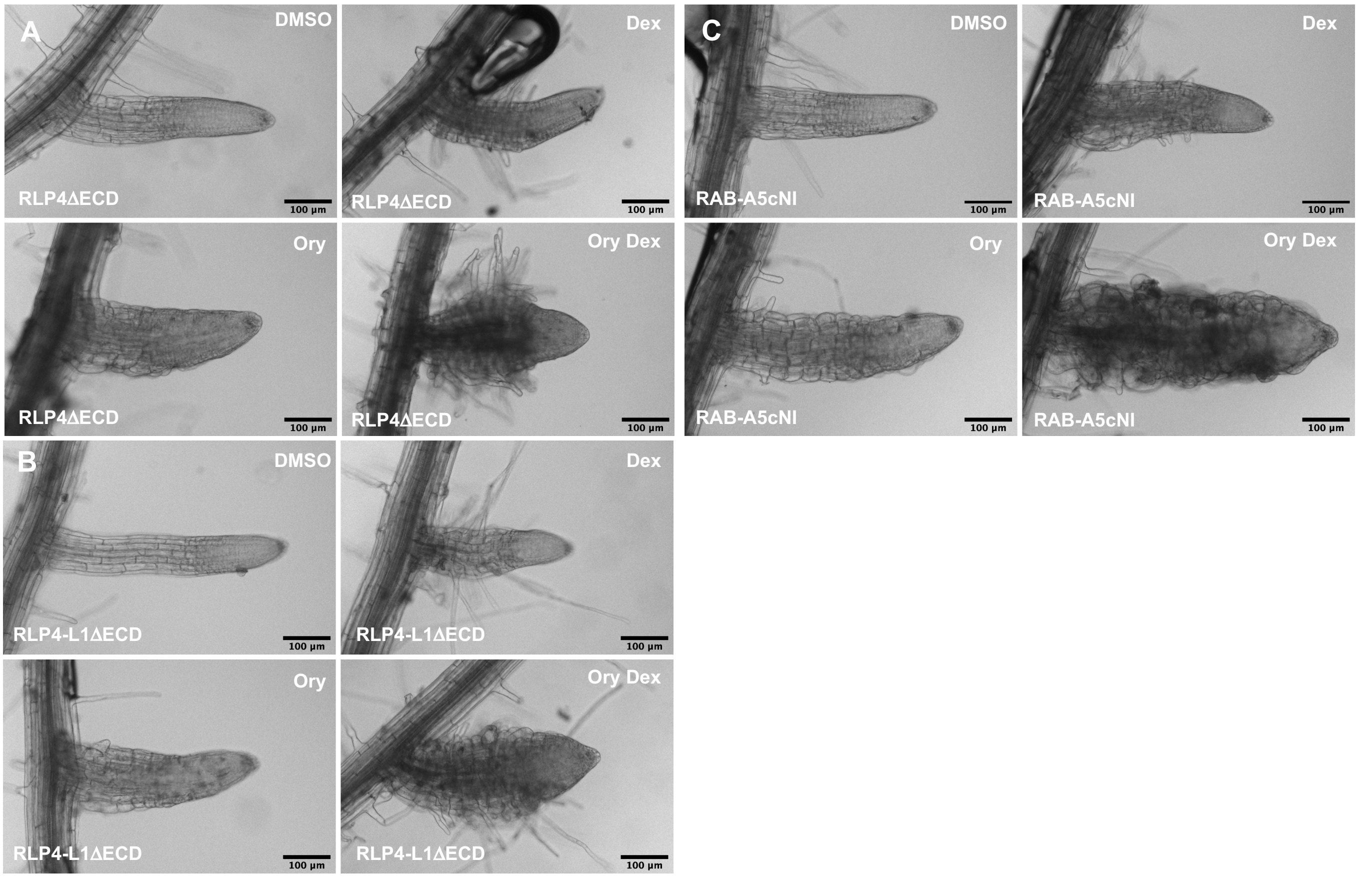


**Figure S5 (related to Figure 5): Inducible over-expression of RLP4s truncated variants causes directional growth defects and hypersensitivity to microtubule inhibition. (A-C)** Photographs of lateral roots expressing *AtRPS5a>>DEX>>RLP4ΔECD* (A), *AtRPS5a>>DEX>>RLP4-L1ΔECD* (B), *and AtRPS5a>>DEX>>RAB-A5c[N125I]* (C) 3 days after transfer to DMSO, 500nM Dex (A,B) or 1µM Dex (C), 250nM Ory, 500nM Dex + 250nM Ory (A,B), or 1µM Dex + 250nM Ory (C). Scale bars 100µm.
